# POZ Domain Transcription Factor Kaiso is a Downstream Effector of STAT1 for Multiple Myeloma Cell Survival and is also essential for Anoikis Resistance of Metastatic Solid Tumors

**DOI:** 10.1101/2025.09.29.675399

**Authors:** Ramadevi Mutra, Faiha Mundodan, Asha Vallem, Charles H. Lawrie, Carla Sole, Carlos Panizo, Prajakta Ghatage, Anciya VR, Sivakumar Vallabhapurapu

## Abstract

Role of the transcription factor STAT1 in cancer cell survival has remained unclear. Here we document an oncogenic role for STAT1 in Multiple Myeloma (MM), wherein STAT1 regulates Kaiso expression, represses *bmf* and is essential for MM cell survival. Moreover, STAT1 is constitutively phosphorylated in MM cell lines and patient derived primary MM cells and STAT1-depletion resulted in reduced Kaiso levels. Importantly, expression of exogenous Kaiso rescued STAT1-depleted MM cells from apoptosis by reversing elevated BMF levels. Further, several MM cell lines harboring diverse oncogenic mutations rely on Kaiso for their survival revealing an oncogenic addiction to Kaiso in MM. Mechanistically, Kaiso interacts with and recruits HDAC1 to pro-apoptotic gene *bmf* promoter to maintain repressive state. In line with this, depletion of Kaiso or HDAC1 results in elevated BMF levels and apoptosis of MM cells. Kaiso is abundantly expressed in several MM cell lines and patient derived primary MM cells. Interestingly, Kaiso interacting partner p120-catenin levels are very low/undetectable in MM indicating that Kaiso function in MM is p120-independent. While previous reports suggested anoikis promoting role for Kaiso in breast cancer, we show here that *bmf* repression by Kaiso is central to anoikis-resistance of metastatic solid tumors. Collectively, our data indicate that Kaiso is a downstream effector for STAT1 in MM cell survival and that Kaiso mediated *bmf* repression is central to MM cell survival and anoikis resistance of metastatic solid tumors. Targetting Kaiso-mediated *bmf* repression would enable us to develop common drug for MM and metastasis of solid tumors.

## Introduction

Cancer Heterogeneity has been major problem in developing anti-cancer drugs (1–7). Moreover, while various pro-tumorigenic transcription factors were shown to operate in progression of multiple cancer types, it is likely that these factors contribute to different cancers by different mechanisms (8–13). Blocking the oncogenic signaling pathways or complete inhibition of a given pro-tumorigenic transcription factor would impair tumor growth, but are often associated with severe toxic side affects due to normal homeostatic requirements of such signaling pathways / transcription factors. Hence, identification of gene-regulatory complexes downstream to oncogenic signals, which regulate commonly operated survival mechanisms within and across different cancers would enable us to develop broad-spectrum anti-cancer drugs by methods disrupting such complexes. To this end, we focussed on the Jak-STAT pathway for novel downstream effectors of this pathway in the context of Multiple Myeloma (MM) and investigated a transcription factor Kaiso for its role in MM cell survival and anoikis resistance of metastatic solid tumors.

Multiple Myeloma (MM) is a cancer of germinal center derived monoclonal plasma cells that has been subdivided into multiple genetic subgroups based on diverse genetic abnormalities (14–22). Several of them have been classified into three major groups of chromosomal translocations involving Cyclin D (Group-1), MAF (Group-2) or MMSET/FGFR3 (Group-3) (19). Although what triggers the malignant transformation of the plasma cells has largely remained unclear, often MM is preceded by a condition known as Monoclonal Gammopathy of Undetermined Significance (MGUS) and in some cases, MGUS cells were known to progress to a state called smouldering myeloma, which is a much milder disease (19). While some of the currently employed therapies have been helpful in MM treatment, it largely remained incurable (23) perhaps due to the heterogeneity of oncogenic mutations across different MM subgroups. Hence, identification of molecular requirements and the survival signals that are common to all genetic subgroups of MM would help develop novel therapeutic approaches to target all MM subgroups.

Among the known oncogenic pathways in MM, the Jak-STAT pathway driven especially by IL-6 to activate STAT3 plays key roles in MM cell survival and cancer progression (19, 24–31). However, the roles of other STATs in MM, especially STAT1 remained elusive. The Jak-STAT pathway is activated by ligand binding to various cytokine receptors that results in the activation of the JAK kinases (32–34). There are four JAK kinases including JAK1, JAK2, JAK3 and TYK2 and different cytokine receptors rely on different JAK kinases to activate downstream STAT members (32, 34). Among the diverse cytokines known to activate the Jak-STAT pathway, Type-I and Type-II interferon signaling pathways activateSTAT1 (35). However, whether STAT1 acts as a tumor promoter or tumor suppressor remained unresolved as opposing roles for STAT1 in different cancers have been reported (36–42). In MM, STAT1 has been suggested to play a role in glycolysis and that it was dispensable for cell survival (43). This is confusing, as glycolysis is linked to cell survival in various cancers (44–47). Further, it was suggested that STAT1 interferes with IL-6 induced STAT3 activity without affecting cell survival in MM (42). Moreover, few other reports suggested that STAT1 activation is linked to MM cell death and inhibition of MM growth based on correlative data suggesting that STAT1 might function as tumor suppressor in MM (48, 49). Likewise, IFN-γ, that activates STAT1 has been shown to act both as tumor promoter and tumor suppressor in different cancers (50–54). While several studies point towards a tumor suppressive role for IFN-γ, some studies have comprehensively revealed a pro-tumorigenic role for IFN-γ. For instance, melanomagenesis caused by UV radiation has been shown to be promoted by IFN-γ but not by type-I Interferons (50). In MM, IFN-γ has been shown to induce apoptosis in MM (49, 55) and has also been shown promote MM growth by inducing PD-L1 and BCL-6 in a STAT1 dependent manner (56, 57). These reports are confusing and convincing evidence is needed to reach a conclusion on whether STAT1 acts as tumor promoter or tumor suppressor in MM. Especially, whether STAT1 plays a role MM cell survival has not been addressed. Collectively, the role of both IFN-γ and STAT1 in MM cell survival and cancer progression has remained enigmatic.

The transcription factor Kaiso (encoded by *ZBTB33*) belongs to the BTB-POZ family of proteins that are characterized by the presence of a N-terminal BTB / POZ domain (Broad complex, Tramtrack and Bric à Brac / Poxvirus and Zinc Finger), wherein the POZ domain mediates protein-protein interactions and the Zinc Finger motif/s contributes to DNA binding and transcriptional regulation (58–62). While the BTB / POZ domain proteins were first found in drosophila (59–61), several of them were subsequently found in humans and other mammals and some of them including B-cell lymphoma-6 (Bcl-6), osteoclast derived zinc finger protein (OCZF), Leukemia/lymphoma related factor (LRF), Fanconi anemia zinc finger (FAZF) protein, hypermethylated in cancer 1 (HIC-1), and testis zinc finger protein (TZFP) were reported to play pro-tumorigenic role (59). Kaiso, however, has been shown to act both as tumor promoter and tumor suppressor (58) and its role in hematological malignancies especially in MM has remained elusive. Importantly, while most POZ-ZF proteins were shown to act as transcriptional repressors, Kaiso has been shown to act both as transcriptional activator and transcriptional repressor (58). Although, initial studies suggested Kaiso to be a transcriptional repressor, ChIP analysis found that Kaiso is largely bound to transcriptionally active regions of chromatin suggesting that it predominantly acts as a transcriptional activator (58, 63). While the mechanistic aspects of transcriptional activation / repression by Kaiso remained largely unanswered, whether Kaiso promoters cancer by acting as a transcriptional activator or repressor has not been clear.

Some of the elegant studies from Juliet M Daniel and Akeel Baig groups have shown that Kaiso gets recruited to methylated DNA sequences to repress target genes or to a Kaiso Binding Sequence (KBS – TCCTGCNA) to activate transcription of target genes (58, 64). However, whether KBS sequence binding of Kaiso would always result in transcriptional activation or whether its KBS binding cause repression of tumor suppressor genes remains to be tested. With regards to the functional regulation of Kaiso, it was first identified as a p120-catenin interacting protein (58) and was thought to participate in p120-associated functions such as regulation of E-Cadherin, cell-cell adhesion, regulation of RhoA, Rac, and Cdc42 as well as actin dynamics (58, 65). Being a transcription factor, while Kaiso has largely been shown to be localized to nucleus, perhaps owing to its interaction with p120, Kaiso has also been shown to be cytoplasmic in solid tumors (58, 66). Moreover, while the physiological importance of p120-mediated Kaiso regulation has remained unclear, under certain conditions, p120 was shown to localize to nucleus, interact with Kaiso and relieve some genes from Kaiso-mediated repression (67, 68). Such p120-regulation of Kaiso has been thought to play key role in cancer progression (69, 70). Importantly, while p120-Kaiso axis has been implicated in solid tumors, whether p120 plays a role in MM and whether Kaiso function is under p120 control in MM has remained unaddressed.

One of the important aspects of tumor metastasis is anoikis resistance (71) which is achieved by active repression of anoikis inducing pro-apoptotic genes *bim* and *bmf* (72–74). While some repressors of *bim* and *bmf* were reported (75, 76), mechanism of their repression exclusively in detached (circulatory) metastatic cancer cells remained unclear. Interestingly, while Kaiso has been implicated in metastasis of solid tumors (58), it was reported to repress *wnt11*, a gene which is required for anoikis resistance of detached breast cancer cells (67). These findings were puzzling and raises a question whether Kaiso promotes anoikis or contributes to anoikis resistance in metastatic cancers.

Collectively, in this study we addressed the critical questions mentioned above and found that STAT1 is oncogenic for MM and Kaiso is a novel downstream effector of oncogenic STAT1 in MM cell survival. Importantly, Kaiso-mediated repression of *bmf* is a commonly operated mechanism in MM cell survival and anoikis resistance of metastatic solid tumors.

## Methods

### Materials and Methods

#### Cell Culture and Cell lines

All Multiple Myeloma Cell lines used in this study were kindly provided by Prof. Leif Bergsagel Lab (Mayo Clinic)and Prof. Marta Chesi (Mayo Clinic). HEK 293T, MDA-MB-231, A-172, HT-29 were purchased from ATCC. MMCL’s were cultured in RPMI-1640 medium (Gibco) Supplemented with with10% FBS (Sigma), 1% Penicillin-Streptomycin (Gibco). Other cell lines were Cultured in DMEM high Glucose Medium (Gibco) supplemented with 10% FBS (Sigma) and 1% Penicillin-Streptomycin (Gibco). All the cell lines were grown in a CO2 incubator maintained with 5% CO_2,_ 37^0^C temperature and 95% humidity.

#### Lentiviral Transduction and Gene Silencing

Pre-designed ShRNA clones Specific to various target genes were procured from Merck-Sigma Aldrich. RNAi Consortium (TRC) developed and provided all the TRCN clones information in the Sigma Aldrich website. TRCN clones specific to ZBTB33 are TRCN0000017838 and TRCN0000232207, TRCN clones specific to STAT1 are TRCN0000004265 and TRCN0000004266, TRCN clones that targets HDAC1 are TRCN0000004816 and TRCN0000197198, TRCN clone that targets STAT2 is TRCN0000364400. pLKO.1-puro-ShNT plasmid which does not target any gene but express a scrambled Sh-RNA was purchased from Addgene (#109012).

Lentiviruses that express different Sh-RNAs or control-ShRNA were prepared by co-transfecting HEK293T cells with the Packaging vectors; Gag, Pol expressing psPAX2 (addgene #12260) and envelope plasmid pMD2.G (addgene # 12259) along with pLKO.1 vector containing Specific gene Targeting Sh RNA. pLKO.1-puro-ShNT was used in place of gene targeting pLKO vector as a control. 72 hours post transfection lentivirus containing supernatant was collected by passing the supernatant through 0.45 µ PES syringe filters. The collected lentivirus was used to transduce either MMCL’s or other solid tumor cell lines in order to silence a target gene. Immunoblotting was performed to check the knockdown efficiency.

#### Plasmids and Transfection

FUW-Kaiso-Flag, FUW-HDAC1-HA, FUW-EGFP were generated in the lab. pGL2-Basic Vector(#E1641) and pSV-β-Galactosidase Control Vector (#E1081) were purchased from Promega. FUW (#14882), pLKO.1-Puro (#8453), pLKO.1-puro-shNT (#109012) were procured from Addgene. FUW-Kaiso-Flag plasmid was generated by cloning Kaiso cDNA with C terminus Flag in to Xba1 and Asc1 sites of FUW vector. FUW-HDAC1-HA vector was generated by subcloning HDAC1 cDNA with C terminus HA tag from HDAC1 Flag (Addgene #13820) in to Xba1 and EcoR1 sites of FUW vector. FUW-EGFP plasmid was generated by subcloning EGFP cDNA from FUGW (Addgene #14883) into Xba1 and EcoR1 sites of FUW vector.

Kaiso Rescue construct was generated in the lab by cloning an expression cassette containing (Ubiquitin Promoter-Human Kaiso cDNA and WPRE-Woodchuck Hepatitis Virus Posttranscriptional Regulatory Element) in to AscI and BamHI sites of pLKO.1 vector expressing Sh-RNA targeting 3’ UTR of Kaiso (TRCN0000017838).

Kaiso expression cassette (Ubiquitin Promoter-Human Kaiso cDNA and WPRE-Woodchuck Hepatitis Virus Posttranscriptional Regulatory Element) was cloned in to AscI and BamHI sites of pLKO.1 vector expressing Sh-RNA targeting STAT1 cDNA (TRCN0000004265). As a control vector EGFP expression cassette (Ubiquitin

Promoter-EGFP and WPRE-Woodchuck Hepatitis Virus Posttranscriptional Regulatory Element) was also cloned in to AscI and BamHI sites of pLKO.1 vector expressing Sh-RNA targeting STAT1 cDNA (TRCN0000004265).

HDAC1 Expression cassette (Ubiquitin Promoter-Human HDAC1 cDNA and WPRE-Woodchuck Hepatitis Virus Posttranscriptional Regulatory Element) was cloned in to AscI and BamHI sites of pLKO.1 vector expressing Sh-RNA targeting 3’ UTR of Kaiso (TRCN0000017838).

Lipofectamine 2000 (Invitrogen) was used to perform Transfections in HEK-293T cells.

#### Cell Viability Assay

Cells were stained with Annexin V PE and 7-AAD using BD Pharmingen Kit and cell viability was analysed by FACS Celesta (BD Biosciences) Flow cytometry. Staining protocol was followed according to manufacturer’s recommendations

#### Xenograft Tumour Growth

Xenograft experiments were carried out at Central Animal House facility, CDFD Hyderabad, as per CDFD Animal ethical committee guidelines. JJN-3 and KMS-26 cells were used to conduct MM Xenograft Tumor growth in nude mice for. 3×10^6^ JJN3 or KMS-26 cells expressing Control-Sh RNA or Sh-RNA targeting a Kaiso or STAT1 were injected subcutaneously in to the anesthetized nude mice along with Matrigel. Cells expressing control-ShRNA were injected on the left side and cells expressing Sh-RNA targeting kaiso or STAT1 were injected on the right side. Prior to injection in mice, cell viability was analyzed by flow-cytometry and found to be equal between control and Kaiso-depleted or STAT1-depleted cells.

#### Immunoblot and quantitative PCR Methods

Whole cell lysates were made in Triton-X 100 lysis buffer (composition-Boston Bioproducts) containing Complete Mini Protease and Phosphatase inhibitors (Roche). For Immunoprecipitations, cells were lysed in IP lysis buffer (Pierce-Thermos Scientific) and then lysates were incubated overnight with indicated antibodies at 4^0^C.Then protein G Dynabeads were added to the overnight incubated antibody and lysate solutions. After 1 hour incubation at 4^0^C beads were washed extensively with IP lysis buffer and the proteins were eluted. Nuclear and Cytoplasmic extracts were prepared as described previously (77). Whole cell lysates, immunoprecipitates and nuclear/cytosolic extracts were analysed by immunoblotting with antibodies to Kaiso (Santacruz,# sc-365428), STAT1 (Cell Signaling Technology # 14994), p-STAT1(Cell Signaling Technology #9167), HDAC1(Cell Signaling Technology #34589), BMF(Cell Signaling Technology #50542), FLAG (Sigma #F3165), HA (Santacruz,# sc-7392), GFP(Santacruz,#sc-9996), β-Tubulin (Cell Signaling Technology #86298), β-Actin(Cloud clone #MAB340Mi21), c-PARP(Cell Signaling Technology #32563), Acetyl histone H3(K9)(Cell Signaling Technology # 9649), LDHA (Cell Signaling Technology, #3582), Mouse IgG Isotype (Santacruz,# sc-2025), Rabbit IgG Isotype (Cell Signaling Technology, #2729), STAT2 (Cell Signaling Technology, #72604), Cyclin D2 (Cell Signaling Technology, #3741), Cyclin D3 (Cell Signaling Technology, #2936), Cyclin E1 (Cell Signaling Technology, # 4129), Cyclin E2 (Cell Signaling Technology, # 4132), Anti-Rabbit IgG, HRP-linked Antibody (Cell Signaling Technology, # 7074), Anti-Mouse IgG, HRP-linked Antibody (Cell Signaling Technology, # 7076). All Primary antibodies were diluted with 1:1000 dilution whereas secondary antibodies were diluted with 1: 4000 dilutions. Immunoblots were developed by using biorad ECL substrates (#170-5061 & 170-5062). Total RNA was isolated using RNeasy Plus kit (Qiagen) for quantitative qRT-PCR analysis of indicated genes. Total RNA was converted to First Strand cDNA with the help of iScript cDNA synthesis kit (Bio-Rad) and quantitative PCR was performed on 96 well CFX Connect RT PCR machine (Biorad).

Primers used were

HDAC1

Forward: 5’CTTCCCCAACCCCTCAGATT3’

Reverse: 5’AGACCTGGCACCCTTTATGG3’

STAT1

Forward: 5’CTGACTTCCATGCGGTTGAA3’

Reverse: 5’AGGGCCATCAAGTTCCATTG3’

BMF

Forward: 5’GAGGTACAGATTGCCCGAAA-3’

Reverse: 5’-CCCCGTTCCTGTTCTCTTCT-3’

β-Actin

Forward: 5’CACCAACTGGGACGACAT-3’

Reverse: 5’ACAGCCTGGATAGCAACG-3’

Bim

Forward: 5’TCAACACAAACCCCAAGTCC-3’

Reverse: 5’TAACCATTCGTGGGTGGTCT-3’

Puma

Forward: 5’CCTGGAGGGTCCTGTACAATCT-3’

Reverse: 5’TCTGTGGCCCCTGGGTAAG-3’

Noxa

Forward: 5’AGCTGGAAGTCGAGTGTGCT-3’

Reverse: 5’TCCTGAGCAGAAGAGTTTGGA-3’

Bax

Forward: 5’GGTTGTCGCCCTTTTTCTA-3’

Reverse: 5’CGGAGGAAGTCCAATGTC-3’

Bak

Forward: 5’GTAGCCCAGGACACAGAGGA-3’

Reverse: 5’ATAGCGTCGGTTGATGTCGT-3’

Bcl2

Forward: 5’GGAGGATTGTGGCCTTCTTT-3’

Reverse: 5’CATCCCAGCCTCCGTTATC-3’

BclW

Forward: 5’GGCGCACCTTCTCTGATCT3’,

Reverse: 5’ATCCACTCCTGCACTTGTCC-3’

BclXL

Forward: 5’-ATGGCAGCAGTAAAGACCA-3’

Reverse: 5’-TCCCGGAAGAGTTCATTCAC-3’

Survivin

Forward: 5’-CCAGATGACGACCCCATAGA-3’

Reverse: 5’-GCACTTTCTTCGCAGTTTCC-3’

For specific amplification of endogenous Vs exogenous Kaiso, the following primer pairs were used: Endogenous 3’-UTR specific: Forward: 5’-TCTGCCCACAAAGGAGACTT-3’ Reverse: 5’ TGGGAGTCAGATGTGTTGGA-3’ and Exogenous Kaiso cDNA specific: Forward: 5’-TTCCCCTTCCATGTTAGCAC-3’ Reverse: 5’ GTTGCTGTTCGCTTGGACTA-3’.

#### Luciferase Assay

Human 2.5 kb BMF gene promoter was amplified using the primers Forward: 5’-GATGGTACCAGGATCACTTGAGCCTG-3’ and Reverse:5’-GAAGCTAGCAAAATACGCCTGCTCG-3’. It was cloned upstream of Luciferase reporter gene using Kpn1 and Nhe1 sites in pGL2-Basic vector resulting in the generation of recombinant BMF-pGL2 plasmid. HEK-293T cells were transfected with BMF-pGL2 plasmid either with FUW or FUW-Kaiso-Flag plasmid. In both conditions pSV-β-Galactosidase Control Vector (Promega # E1081) was co-transfected. After 36 hours of transfection, lysates were made in 1X Reporter lysis buffer (Promega Cat# E397A) containing protease inhibitor cocktail. Equal Concentration of lysates were used to perform Luciferase assay upon incubating with Luciferase assay substrate (Promega Cat# E151A) with the help of Promega Luminometer. β-gal assay was performed with 2-Nitrophenyl β-D-galactopyranoside (Sigma Cat#: N1127). β-gal activity has been measured as internal control for transfection efficiency. Upon normalising against β-gal values, promoter activity has been plotted as fold change in relative light units (RLU).

#### Chromatin Immunoprecipitation Assay

ChIP was performed as per the manufacturer protocol (Active Motif) with minor modifications whereever required as follows. Briefly myeloma cells were crosslinked with 1% Formaldehyde and cells were subjected to cell lysis and sonication in Sonication buffer with 25% amplitude, 20sec on and 30sec off until we got chromatin of 300-500bp (checked by reverse crosslinking and agarose gel electrophoresis). Sheared chromatin was incubated with indicated antibodies overnight in ChIP buffer 1 containing Protease inhibitor cocktail at 4^0^C. Protein G Magnetic Beads were blocked with 25µl of salmon sperm DNA (1MG/ML-Invitrogen) and 10µl of BSA (2.5MG/ML Sigma) for an hour at 4^0^C, resuspended in chip buffer I and added to the ChIP reaction mix and incubated for 3hours at 4^0^C. Finally, beads were separated by magnetic separator, were washed once with chip buffer 1 and twice with chip buffer2 and eluted with elution buffer provided in the kit. To the eluted chromatin was incubated overnight at 65°C with reverse crosslinking buffer and subjected to Proteinase K treatment. Then the DNA was purified using PCR cleanup columns. PCR was performed by using specific primers designed for region of BMF promoter (−2139 to −1943) Harboring the Kaiso binding sequence (KBS) or a downstream region without KBS (−1475 to −1250) Kaiso Binding Sequence (KBS-TCCTGCNA / TNGCAGGA).

#### Anoikis Assay

60 hours post-transduction with lentiviruses expressing control-Sh RNA or Kaiso-ShRNA, cells were trypsinized and seeded them either in adherent plates or ultra-low adhesion plates to keep them in suspension. After culturing cells in suspension or adherent conditions for 24 hours whole cell lysates were made and were analysed by immunoblotting. To investigate relative anoikis resistance in Control-Sh RNA Vs Kaiso-Sh RNA cells, 24 hours after culturing in suspension conditions, equal number of cells were re-seeded in adherent plates and were further cultured under adherent conditions for another 24 hours. Cells were washed and stained with Crystal violet. Plates containing Crystal violet-stained cells were dried completely at RT for a day and then the bound dye was eluted using 10% acetic acid. Absorbance was measured for eluted dye in multimode plate reader at 595nm. The results were plotted in a bar diagram to show relative survival of the cells to check the anoikis resistance.

## Results

### Oncogenic addition to Kaiso in multiple myeloma

Role of Kaiso in hematological malignancies especially in MM has not been addressed. We show here that Kaiso is abundantly expressed in various MM cell lines and primary patient derived MM cells (Fig. 1 A and B). To find whether Kaiso is essential for MM cell survival, we employed Sh-RNA mediated silencing of Kaiso and found that several MMCLs were subjected to apoptosis upon Kaiso-depletion (Fig. 1C, Supplementary Fig. 1A). Importantly, our results identified an oncogenic addiction to Kaiso in MM as different genetic subgroups of MM cells harboring diverse oncogenic mutations were found to rely on Kaiso for their survival (Fig. 1C). In line with the requirement of Kaiso for MM cell survival in vitro, Kaiso-depletion has completely abrogated human MM tumor growth in vivo in nude mice (Fig. 1D). To rule out off-target effects of Kaiso-ShRNA on MM cell apoptosis, we infected JJN3 cells with a lentiviral vector that simultaneously expresses sh-RNA targeting the 3′-UTR of endogenous Kaiso mRNA and exogenous Kaiso cDNA that is resistant to Kaiso specific sh-RNA (Fig. 1E and Supplementary Fig. 1B). As shown in Fig. 1F, presence of exogenous sh-resistant Kaiso completely rescued JJN3 cells from apoptosis induced by depletion of endogenous Kaiso revealing essential role of Kaiso in MM cell survival.

**Figure 1:**
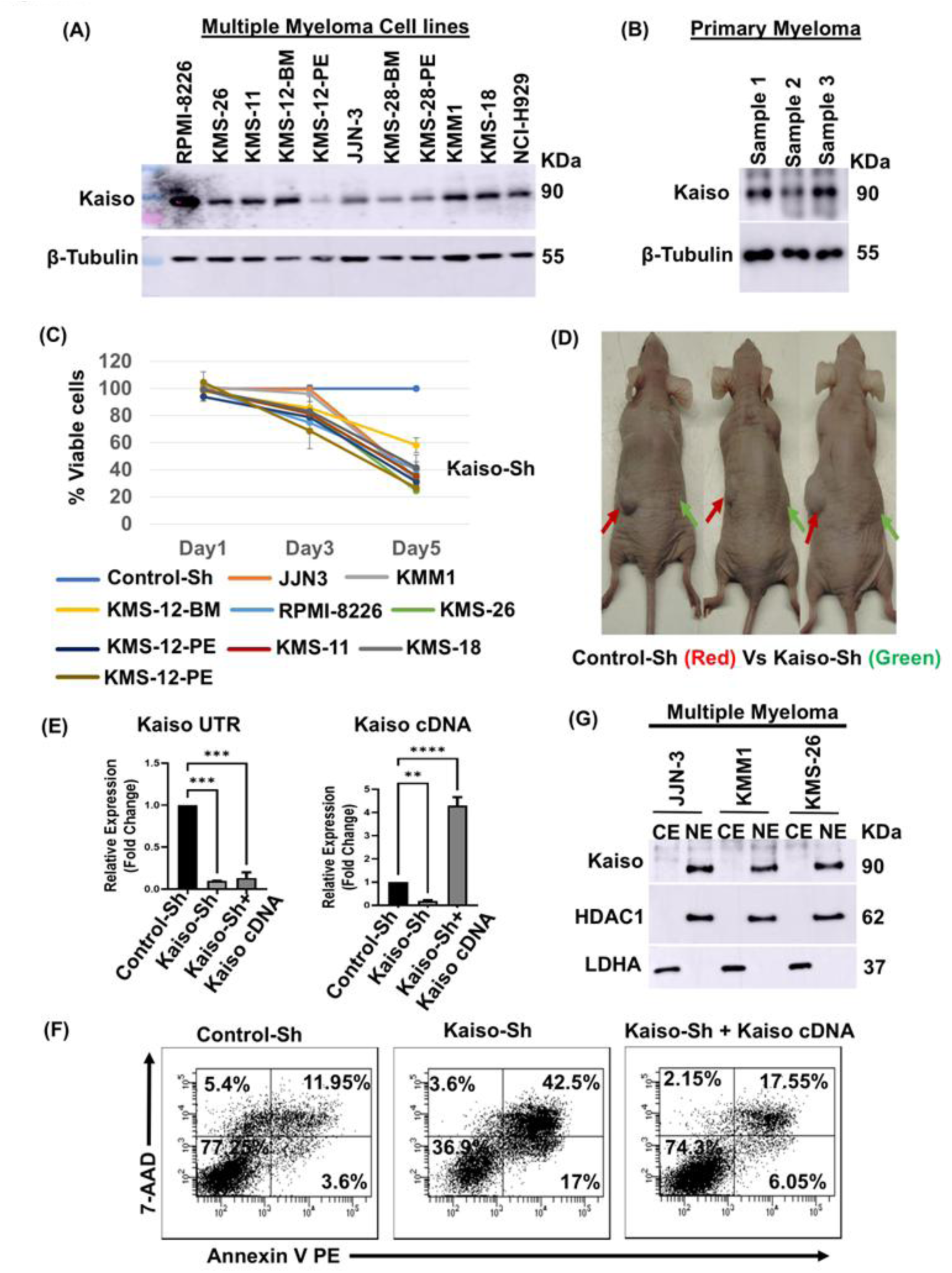
Oncogenic addiction to Kaiso in Multiple Myeloma. (A) Whole cell lysates from the indicated myeloma cell lines and (B) Primary Patient derived Myeloma cells were analysed for Kaiso expression by immunoblotting. (C) Indicated Myeloma cells were transduced with lentiviruses expressing Control-ShRNA or Kaiso-ShRNA. Transduced cells were collected at different time intervals and their survival was analyzed by flow-cytometry upon staining with Annexin V-PE and 7-AAD and the data was presented in the form of a line graph. For convenience, cell viability in control cells was converted to 100% and the viability in Kaiso-depleted cells was normalized accordingly. (D) JJN-3 cells were transduced with Lentiviruses expressing Control-ShRNA or Kaiso-ShRNA. 24 hours post infection, equal number of Control and Kaiso-depleted cells (3×10^6^ cells) were subcutaneously injected into nude mice along with Matrigel. Prior to injection in mice, cell viability was analyzed by flow-cytometry and found to be equal between control and Kaiso-depleted cells. Control cells were injected on the left side and indicated with red arrow. Kaiso-depleted cells were injected on the right side indicated with green arrow. Note the complete block in MM tumour growth upon Kaiso depletion. (E) Total RNA from JJN-3 cells transduced with Lentivirus expressing Control-ShRNA or Kaiso-ShRNA or Kaiso Rescue Construct (Kaiso-Sh+ Kaiso-cDNA) was isolated 60 hours post infection and RT-PCR analysis was performed to analyze the relative expression of Endogenous and Exogenously overexpressed Kaiso. (F) JJN-3 cells were transduced with Lentivirus expressing Control-ShRNA or Kaiso-ShRNA or Kaiso Rescue Construct (Kaiso-Sh+ Kaiso-cDNA) and 5 days later cell viability was analysed by flowcytometry upon staining with Annexin V-PE and 7-AAD.(G) Cytosolic Vs Nuclear distribution of Kaiso among different MM cells was analysed by immunoblotting using indicated antibodies.

Importantly, while Kaiso function in solid tumors was shown to be controlled by its interacting partner p120-catenin (p120) (58), we show here that p120 levels were not detectable in MM cells (Supplementary Fig. 1C). Perhaps owing to its interaction with p120, Kaiso was shown to be distributed both in cytoplasm and nucleus in solid tumors (66, 70) (Supplementary Fig. 1D). However, in MM cells, Kaiso was found to be almost completely localized to nucleus (Fig. 1G and Supplementary Fig. 1D) probably due to lack of p120 expression. Importantly, it was previously shown that higher nuclear levels of Kaiso results in more aggressive breast cancer (66). Hence, in MM, Kaiso being restrained to nucleus appear to cause aggressive cancer growth. Collectively, these data suggest that Kaiso function in MM is independent of p120 and that p120 has no role in MM cell survival.

### Kaiso Represses *bmf* in Multiple Myeloma

To investigate the mechanism by which Kaiso regulates MM cell survival, we analyzed the expression of various Bcl-2 family members in Kaiso-depleted cells (Supplementary Fig. 2A). While we did not find significant changes in expression of several anti- or pro-apoptotic Bcl-2 family members (Supplementary Fig. 2B), we repeatedly observed elevated levels of *bmf* upon Kaiso-depletion (Fig. 2A and B and Supplementary Fig. 2D). Similar results were obtained with two different Sh-RNAs specific to Kaiso. We then, investigated how Kaiso represses *bmf*. To this end, we first analyzed a 2.5kb *bmf* promoter region by bio-informatic methods and found potential Kaiso binding sequence (KBS) at −2033 to −2025 (Supplementary Fig. 2C). We then proceeded to clone the 2.5kb *bmf* promoter into pGL2-basic luciferase vector and investigated whether it is regulated by Kaiso. To this end, luciferase reporter assays concluded that Kaiso represses *bmf* promoter (Fig. 2C and Supplementary Fig. 2E). To find whether Kaiso represses *bmf* by directly binding to its promoter in MM cells, we employed chromatin immunoprecipitation (ChIP) experiments and found that Kaiso is recruited to a KBS site located at −2139 to −1943 region but not to a downstream region (−1475 to −1250) within *bmf* promoter suggesting specific Kaiso recruitment to −2139 to −1943 region harboring KBS (Fig. 2D and Supplementary Fig. 2F) Importantly, while Kaiso-depletion resulted in elevated BMF levels, expression of Sh-RNA resistant Kaiso-cDNA completely reversed BMF levels in Kaiso-depleted cells indicating specific requirement of Kaiso for *bmf* repression (Fig. 2E and F). Moreover, in line with the increased apoptosis, cleaved PARP levels also have increased in Kaiso-depleted cells (Supplementary Fig. 2G).

**Figure 2.**
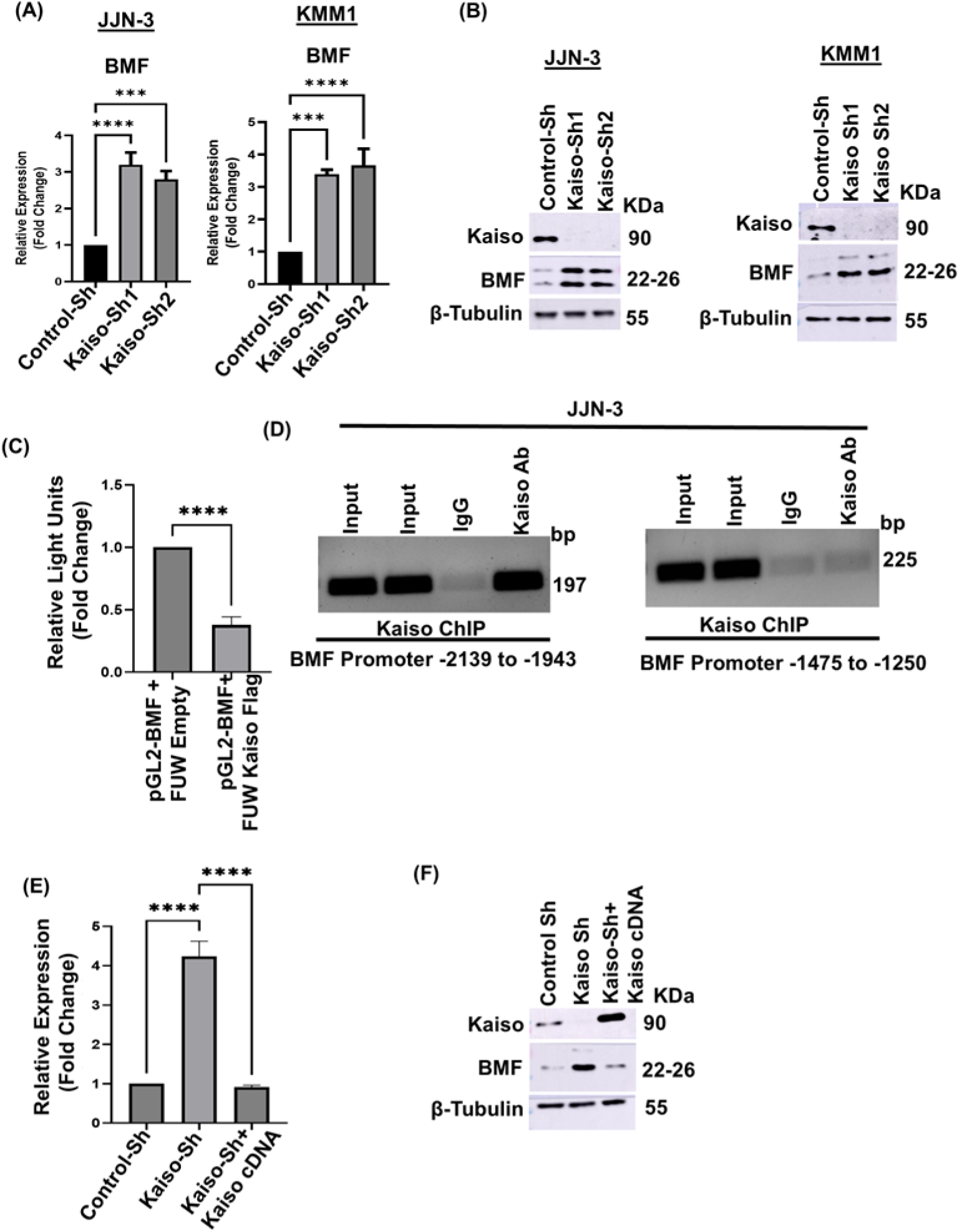
Kaiso Represses BMF gene in Multiple Myeloma: (A) Total RNA from the indicated Control and Kaiso-depleted MM cells was collected and the relative expression of BMF was analysed by real time PCR. JJN3 and KMM1 cells were infected with Lentivirus expressing Control-ShRNA or Kaiso-ShRNA. 60 hours post transfection total RNA and whole cell lysates were made and RT-PCR analysis(A) immunoblotting (B) were performed for the indicated genes/proteins respectively. (C) Human 2.5kb BMF-pGL2 promoter activity was measured by Luciferase reporter assay in HEK-293T cells both in the presence and absence of Kaiso-Flag. (D) Chromatin from JJN3 cells was subjected to immunoprecipitation using anti-Kaiso antibody or normal mouse IgG. Kaiso binding to BMF promoter was analyzed by PCR amplification of chromatin-immunoprecipitated DNA using primers corresponding to a region of BMF promoter (−2139 to −1943) harboring Kaiso binding sequence or a region (−1475 to −1250) lacking Kaiso Binding Sequence. (E) JJN-3 cells were transduced with Lentivirus expressing Control-ShRNA, Kaiso-ShRNA, Kaiso Rescue Construct (Kaiso-ShRNA+ Kaiso-cDNA). 60 hours post transfection, total RNA and whole cell lysates were collected and the relative expression of BMF was analysed by RT-PCR (E) and immunoblotting (F).

Collectively, these results highlight the essential requirement of Kaiso-mediated *bmf* repression in MM cell survival.

### Kaiso Interacts with HDAC1 in MM

To investigate the mechanism by which Kaiso represses *bmf* in MM, we first analyzed with which of the histone modifiers Kaiso interacts and found that it frequently interacts with HDAC1 in MM as revealed by immunoprecipitation experiments (Fig. 3A and B). Also, upon co-transfection in HEK-293T cells Kaiso and HDAC1 were found to interact with each other as shown by immunoprecipitation experiments (Fig. 3C and Supplementary Fig. 3A). We next investigated whether *bmf* is under HDAC mediated repression in different MM cells, we treated JJN3 and NCI-H929 cells with HDAC-inhibitor TSA and found that BMF is significantly elevated (Supplementary Fig. 3B).

**Figure 3:**
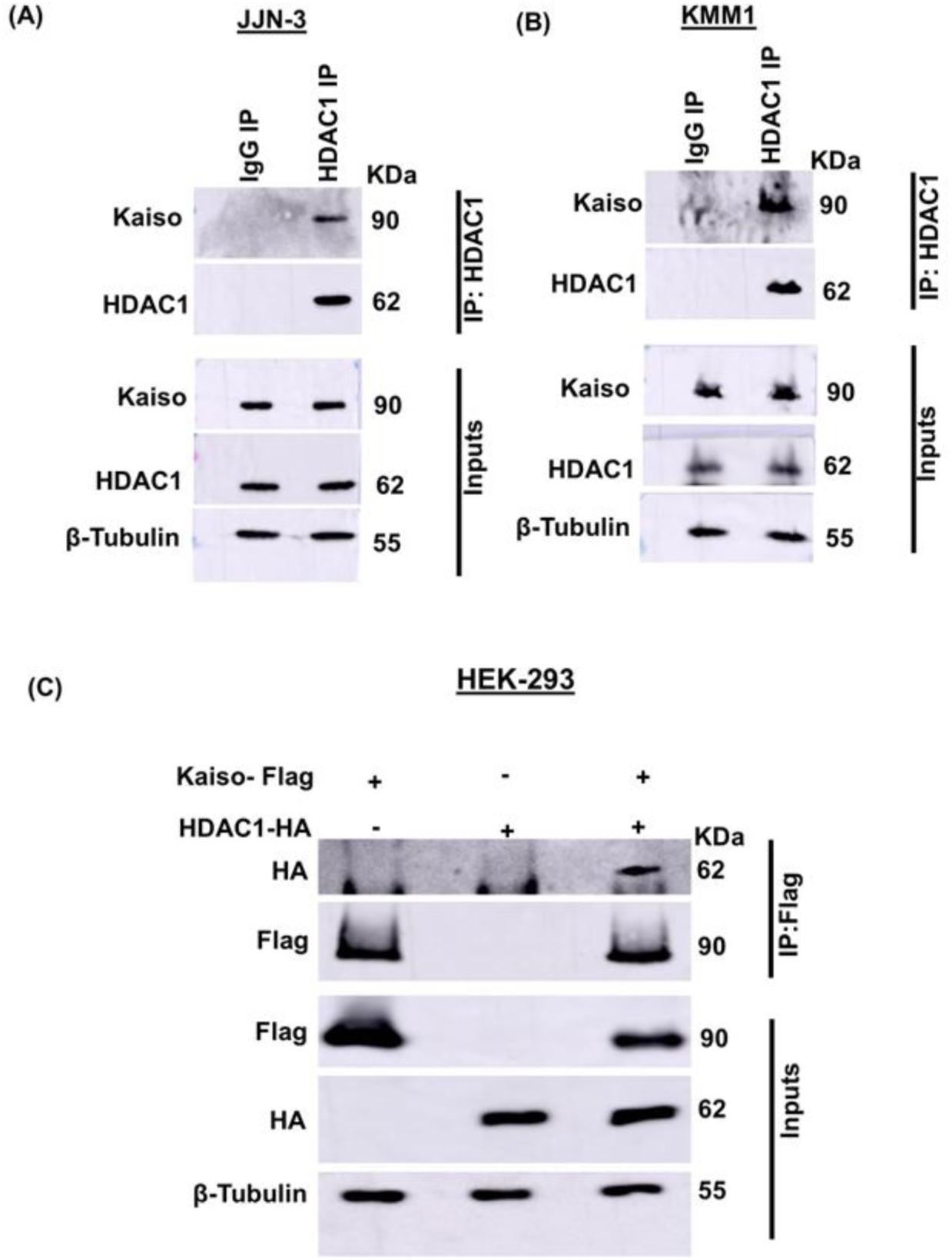
Kaiso interacts with HDAC1 in Multiple Myeloma: Whole cell lysates from JJN3 (A) and KMM1 (B) were subjected to immunoprecipitation using anti-HDAC1 antibody or normal rabbit IgG. Interaction of HDAC1 and Kaiso was analysed by immunoblotting as indicated (C) HEK-293T cells were co-transfected with Kaiso-Flag and HDAC1-HA and immunoprecipitation was performed with anti-Flag antibody. Interaction between HDAC1 and Kaiso was analysed by immunoblotting as indicated.

### Kaiso-HDAC1 complex maintains repressive chromatin on *bmf* gene promoter

While the above results identify the formation of Kaiso-HDAC1 complex in MM, whether HDAC1 is essential for *bmf* repression is not known. To this end, we first analysed the expression of HDAC1 in several MMCLs and found that HDAC1 is abundantly expressed in most MM cells (Supplementary Fig. 4A). We then silenced HDAC1 by employing lentiviral mediated expression of HDAC1-specific Sh-RNAs. Two different HDAC1 specific Sh-RNAs were employed and found that HDAC1-depletion results in significantly elevated BMF RNA and protein levels (Fig. 4A and B). In line with elevated BMF levels, HDAC1-depletion also results in MM cell apoptosis (Fig. 4C). Importantly, HDAC1-depletion did not affect the expression of other HDACs in MM cells (Supplementary Fig. 4B). Next, we investigated whether HDAC1 is recruited to *bmf* gene promoter by ChIP experiments and found that HDAC1 is also recruited to −2139 to −1943 region of *bmf* promoter where Kaiso is recruited (Fig. 4D left panel). Interestingly, HDAC1 recruitment to *bmf* promoter was completely abrogated in Kaiso-depleted cells, indicating that Kaiso recruits HDAC1 to *bmf* promoter (Fig. 4D middle panel). HDAC1 levels were not altered in Kaiso-depleted cells (Supplementary Fig. 4C) suggesting that the loss of HDAC1 recruitment to *bmf* promoter in Kaiso-depleted cells was not due to reduced HDAC1 protein levels. In line with the absence of HDAC1 recruitment to the *bmf* promoter in Kaiso-depleted cells, recruitment of acetylated histone H3 was found to be elevated (Fig. 4D right panel) resulting in increased expression of BMF. We next investigated whether HDAC1 over expression in Kaiso-depleted cells would rescue MM cells from apoptosis and reverse BMF levels. To this end, we made a lentiviral vector that simultaneously express Kaiso-ShRNA and HDAC1 (Supplementary Fig. 4D and Fig. 4E) and found that HDAC1 overexpression does not rescue Kaiso-depleted cells from apoptosis (Fig. 4E). Also, HDAC1 overexpression does not reverse BMF levels in Kaiso-depleted cells (Supplementary Fig. 4E) indicating that HDAC1 strictly depends on Kaiso to repress *bmf*.

**Figure 4:**
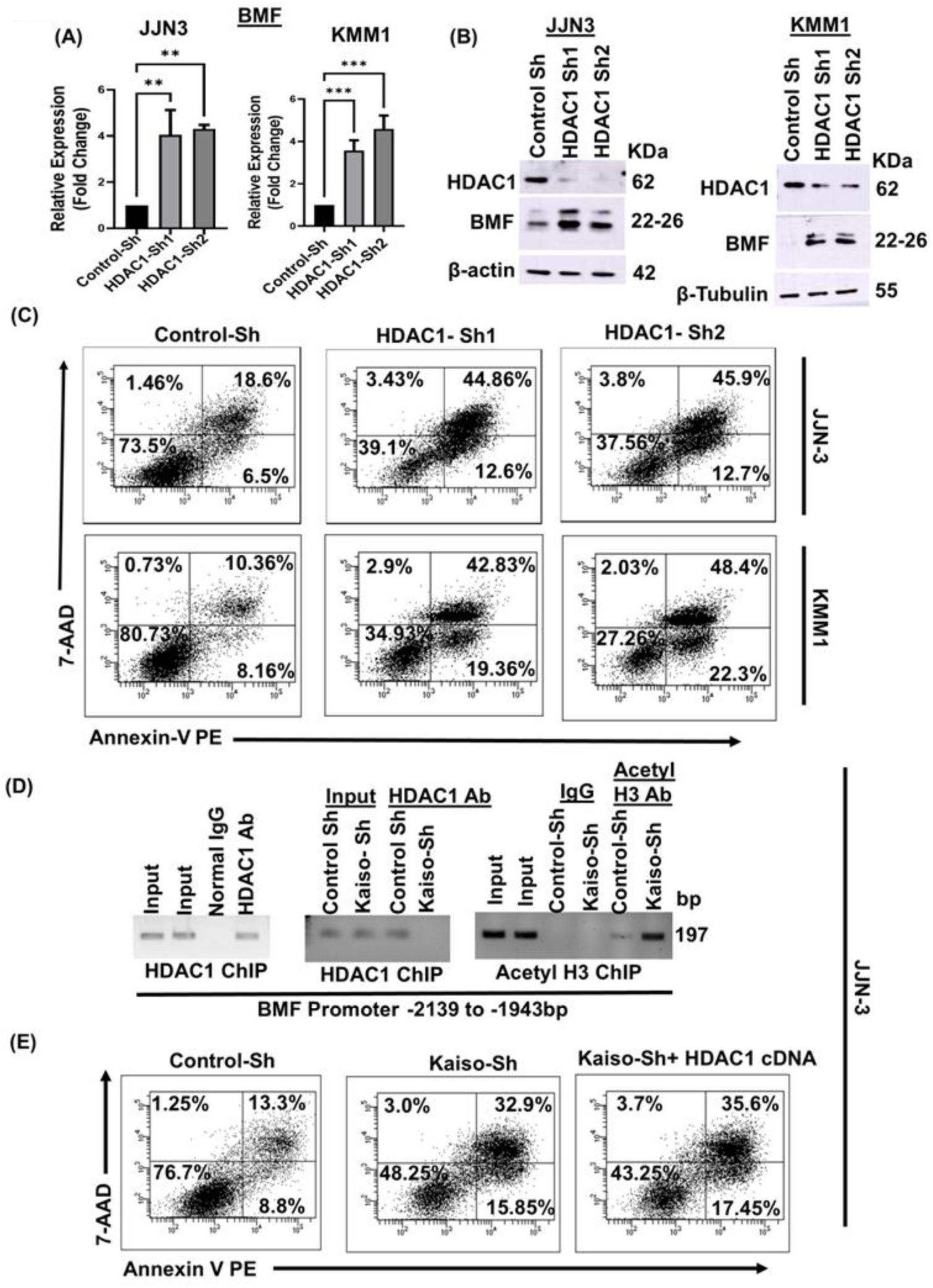
Kaiso-HDAC1 complex maintains repressive chromatin on BMF promoter. (A) HDAC1 was depleted in indicated myeloma cell lines using HDAC1-ShRNA expressing lentivirus. 60 hours post-infection cells were collected to make Total RNA and Whole cell Lysates. (A) RT-PCR analysis was performed to check the relative Expression of BMF (Left JJN3 & Right -KMM1) and (B) immunoblotting for the indicated proteins (Left JJN3 & Right -KMM1). (C) 5 days post transduction, cell viability was analysed by flowcytometry upon staining with Annexin V-PE and 7-AAD (Top panel-JJN-3 & Bottom panel -KMM1). (D) Chromatin from JJN-3 was immunoprecipitated using anti-HDAC1 antibody and HDAC1 binding to BMF promoter was analysed by PCR amplification of immunoprecipitated DNA using primers corresponding to −2139 to −1943 of BMF promoter (Left). Chromatin from Control or Kaiso-depleted JJN3 cells was immunoprecipitated with anti-HDAC1 antibody (Middle) and anti-Acetyl H3 antibody (Right) and HDAC1 binding to BMF promoter or histone acetylation status of BMF Promoter was analysed by PCR amplification of immunoprecipitated DNA using the primers as mentioned above. Normal Rabbit IgG antibody was used as control. (E) JJN3 cells were transduced by lentiviruses expressing control-ShRNA or Kaiso-ShRNA or Kaiso-ShRNA + HDAC1 cDNA. 5 Days post-infection cells were collected and viability was analysed by flowcytometry upon staining with Annexin V-PE and 7-AAD. Note that exogenous expression of HDAC1 was insufficient to rescue Kaiso-depleted cells from apoptosis.

Collectively, these results highlight that Kaiso-HDAC1 complex mediated repression of *bmf* is an essential survival strategy operated in MM.

### STAT1 activated by the Jak-STAT pathway regulates Kaiso Expression in MM

The above results established an important role for Kaiso in MM cell survival and cancer growth. However, upstream signals involved in Kaiso expression in MM are not known. To this end, we investigated different signaling pathways and found that blocking of Jak-STAT pathway results in reduced levels of Kaiso in MM (Fig. 5A). While Jak-STAT pathway activates different STATs, our results revealed that Sh-RNA mediated silencing of STAT1 results in significantly downregulated Kaiso (Fig. 5B) indicating the essential role of STAT1 for Kaiso expression in MM.

**Figure 5:**
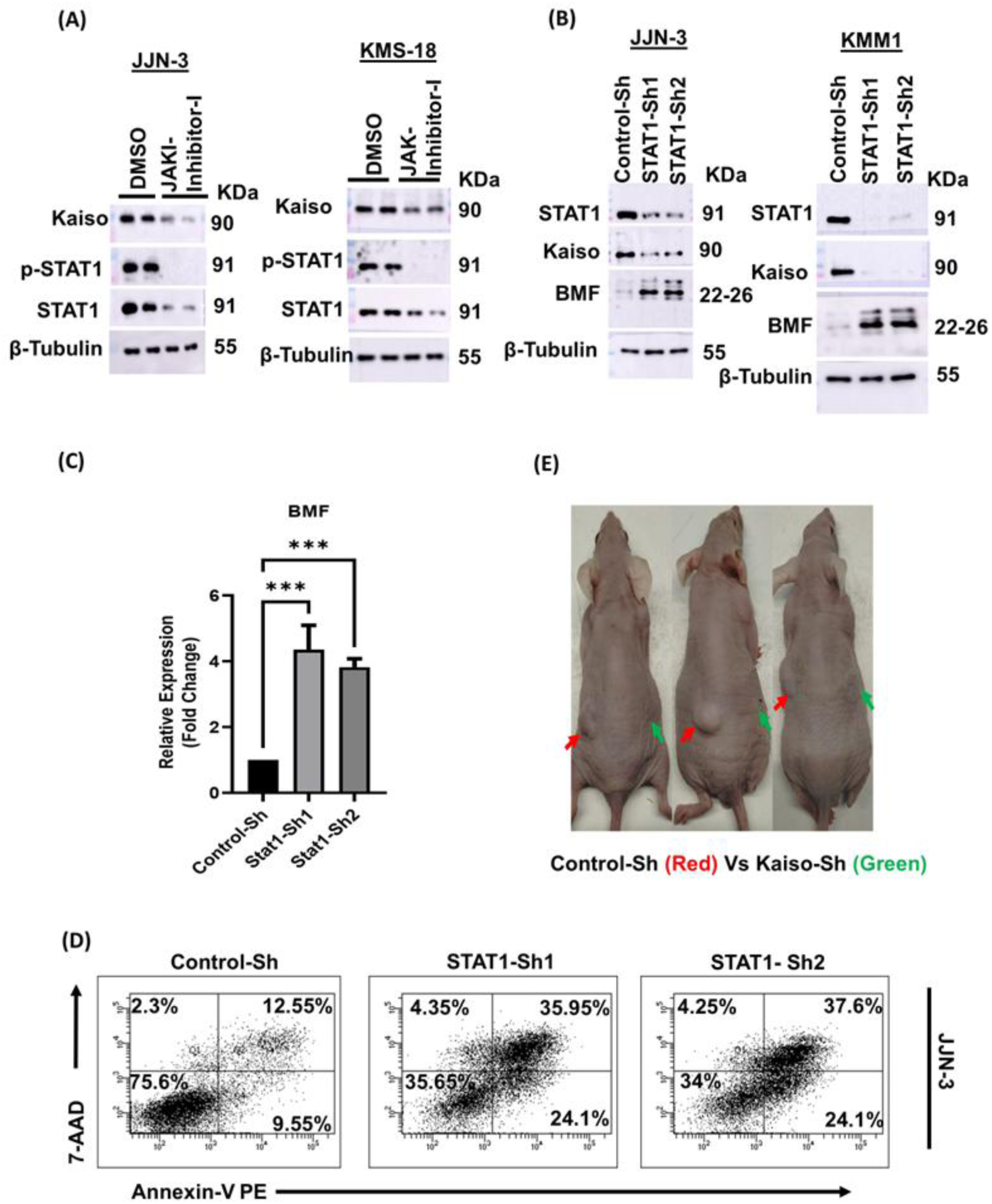
STAT1 is oncogenic in MM and Kaiso is a downstream effector of STAT1 in Multiple Myeloma cell survival. **STAT1 is oncogenic in MM and regulates Kaiso expression.** (A) JJN-3 and KMS-18 cells were treated with Jak-STAT inhibitor-I or DMSO for 8 hours. Whole cell lysates were made and the levels of indicated proteins were analysed by immunoblotting. (B) JJN-3 and KMM1 cells were transduced with lentiviruses expressing Control-ShRNA or STAT1-ShRNA. 60 hours post infection whole cell lysates were prepared and analysed for indicated proteins by immunoblotting and (C) Total RNA from control and STAT1-depleted JJN3 cells was made and RT-PCR analysis was performed for the relative expression of BMF. Note that STAT1 depletion resulted in the significant increase in BMF levels like in the case of Kaiso Knockdown. (D) JJN-3 cells were transduced with Lentiviruses expressing Control-ShRNA or STAT1-ShRNA. 5 days post infection, cell viability was analysed by flow-cytometry upon staining with Annexin V-PE and 7-AAD. (E) KMS-26 cells were transduced by lentiviruses expressing Control-ShRNA or Kaiso-ShRNA. 24 hours post infection, cell viability was found to be equal in both groups of cells and equal number of Control or STAT1-depleted cells (3×10^6^ cells) were subcutaneously injected into nude mice along with Matrigel. Control cells were injected on the left side and indicated with red arrow. STAT1-depleted cells were injected on the right side indicated with green arrow. Note the complete block in Myeloma tumour growth upon STAT1 depletion.

### Oncogenic role of STAT1 in MM

While STAT1 has been largely shown to act as a tumor suppressor, its role as a tumor promoter has also been documented in different cancers (37, 42, 43, 48). However, conclusive evidence for the requirement of STAT1 in MM cell survival and cancer growth has been lacking. Our results above indicated that STAT1 is essential for Kaiso expression. In line with this, silencing of STAT1 subjected MM cells to apoptosis (Fig. 5D and Supplementary Fig. 5A). In line with its requirement for MM cell survival, STAT1-depletion has also strikingly abrogated subcutaneous MM cancer growth in nude mice (Fig. 5E). Interestingly, like in the case of Kaiso-depletion, STAT1-depletion also has resulted in significantly elevated BMF levels (Fig. 5B and 5C). Moreover, STAT-1 specific Sh-RNA did not effect expression of other STAT molecules indicating the specificity of STAT-1 Sh-RNA (Supplementary Fig. 5B). We next investigated whether STAT1 contributes to MM cell survival as a heterodimer with STAT2 or independent of STAT2. To this end, we depleted STAT2 in MM cells and found that STAT2-depletion does not effect expression of Kaiso, STAT1 or BMF and does not cause MM cell apoptosis (Supplementary Fig. 5C and 5D). Thus, oncogenic role of STAT1 in MM is independent of STAT2. In addition to regulation of Kaiso expression and BMF repression, STAT1 also seems necessary for expression of Cyclin D and Cyclin E (Supplementary Fig. 5E and 5F). Collectively, these results indicate an oncogenic role for STAT1 in MM and that STAT1 regulates Kaiso expression in MM.

Because STATs regulate gene expression upon phosphorylation, we tested phosphorylation status of STAT1 (Tyr-701) and found that it is constitutively phosphorylated in several MMCLs and primary patient derived MM cells (Fig. 6A).

**Figure 6:**
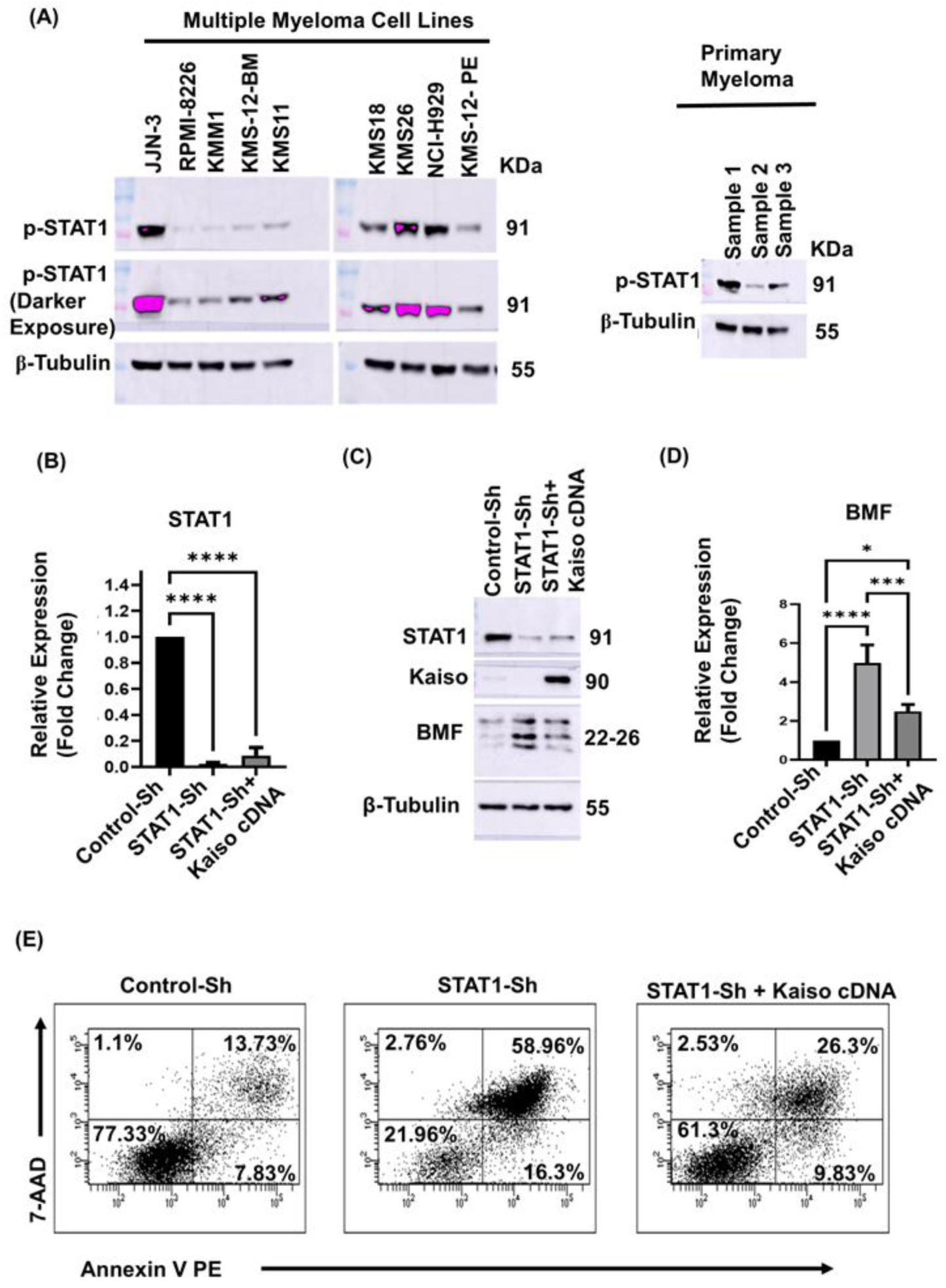
Kaiso is a downstream effector of STAT1 in Multiple Myeloma Cell Survival. (A) Whole cell lysates from the indicated Myeloma cell lines and primary patient derived Myeloma cells were analysed by immunoblotting for the indicated proteins. Note that STAT1 is constitutively phosphorylated most MM cell lines and primary MM cells. (B-D) JJN-3 cells were transduced with lentiviruses expressing Control-ShRNA or STAT1-ShRNA or STAT1-ShRNA+ Kaiso cDNA. 60 hours post-infection total RNA (B&D) and whole cell lysates (C) were collected and analyzed by RT-PCR for the expression levels of indicated genes (B&D) and by immunoblotting for relative levels of indicated proteins (C). (E) JJN3 cells transduced by different lentiviruses as indicated in (B-D) were analysed for cell viability 5 days post infection upon staining with Annexin V-PE and 7-AAD. Note that Exogenous expression of kaiso in STAT1 depleted cells significantly rescued from apoptosis.

### Kaiso is a novel downstream effector of STAT1 in MM cell survival

While the above results indicate that both STAT1 and Kaiso repress *bmf* in MM and that STAT1 regulates Kaiso expression, it was not clear whether Kaiso acts a downstream effector of STAT1 for *bmf* repression and MM cell survival. To this end, we made a lentiviral vector that simultaneously express STAT1-ShRNA and exogenous Kaiso (Fig. 6B and C and Supplementary Fig. 6A). While lentiviral mediated expression of STAT1-ShRNA results in upregulation of BMF and subjects MM cells to apoptosis, expression of exogenous Kaiso in STAT1-depleted cells significantly reversed BMF levels (Fig. 6C and 6D) and rescued STAT1-depleted MM cells from apoptosis (Fig. 6E). As a control, we overexpressed GFP in STAT1-depleted cells and did not observe rescue from apoptosis and BMF levels were not reversed (Supplementary Fig. 6B and 6C).

Collectively, our results identify Kaiso as a novel downstream effector of STAT1 in MM cell survival.

### Kaiso is a key regulator of anoikis resistance in metastatic solid tumors

Normal epithelial cells undergo death when they become non-adherent due to anoikis caused by increased levels of BMF and Bim (72–74). However, metastatic solid tumors exhibit anoikis resistance to be able to metastasize to distant locations (73). Since our results indicated that Kaiso is a key repressor of *bmf* in MM, we asked whether Kaiso also represses *bmf* in metastatic solid tumors and contribute to anoikis resistance. To this end, we depleted Kaiso in metastatic breast cancer (MDA-MB-231), Glioma (A-172) and colorectal cancer (HT-29) cell lines and cultured them either in adherent conditions or non-adherent conditions (ultra-low adherent plates). Interestingly, depletion of Kaiso in breast cancer or glioma cells did not affect BMF levels as long as the cells were cultured under adherent conditions. However, depletion of Kaiso in cells cultured under non-adherent conditions resulted in significant increase in BMF levels (Fig. 7A and Supplementary Fig. 7A). In line with this, Kaiso-depletion did not affect cell viability when cultured in adherent conditions but Kaiso-depletion has resulted in significant loss of cell viability upon culturing in non-adherent conditions (Fig. 7B and Supplementary Fig. 7B). Importantly, Kaiso was recruited to *bmf* promoter both in adherent and non-adherent breast cancer cells (Fig. 7C). These results indicate that Kaiso is essential for the survival of metastatic cancer cells exclusively during the detached phase. In case of colorectal cancer cells (HT-29) however, Kaiso-depletion resulted in moderate increase in BMF levels associated with moderate loss of viability in adherent cells, where as significant increase in BMF levels and enhanced loss of viability was observed in Kaiso-depleted cells cultured in non-adherent suspension condition (Supplementary Fig. 7C and D).

**Figure 7:**
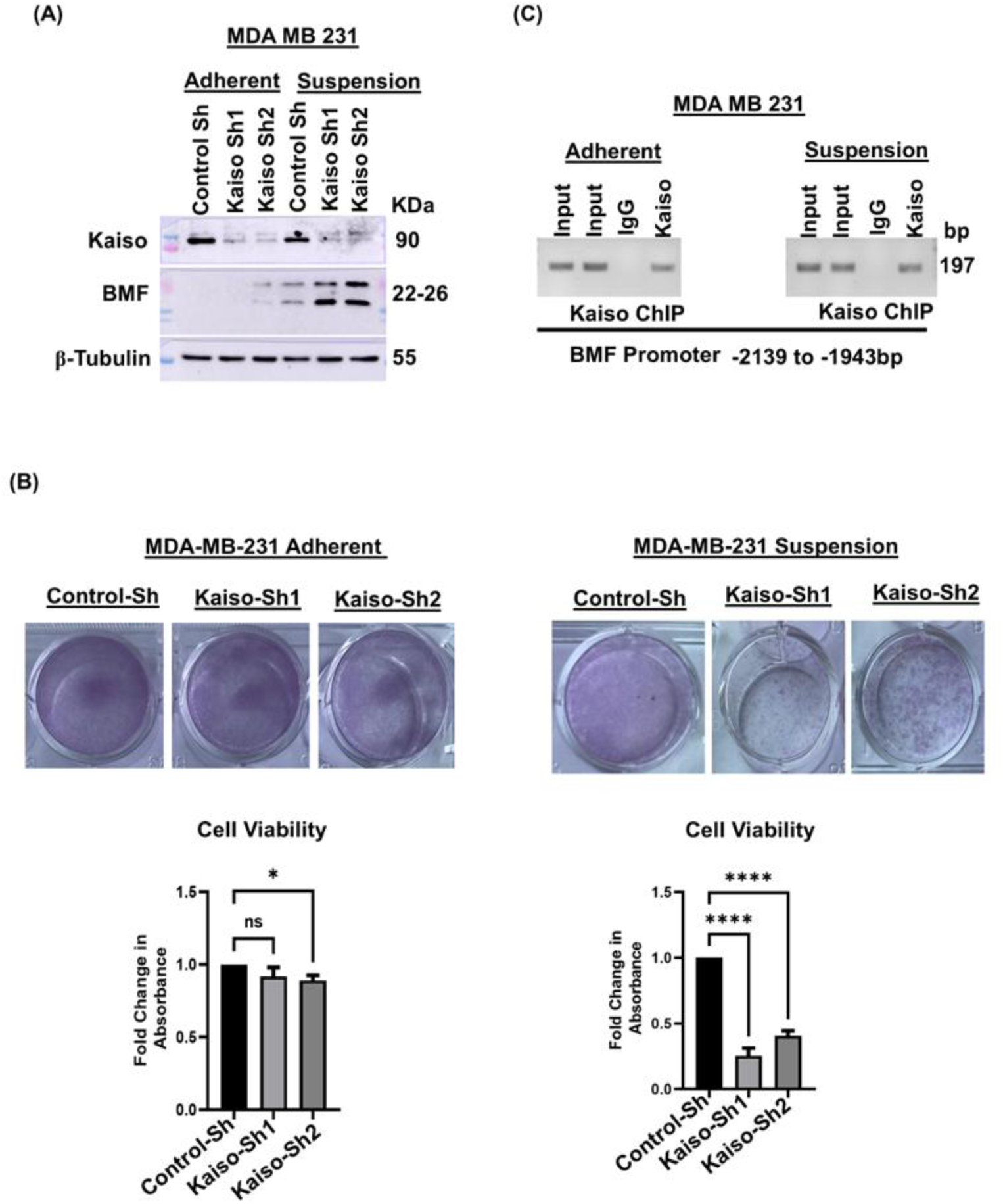
Kaiso is essential for anoikis resistance in Metastatic Solid Tumors. (A) MDA-MB-231 cells were transduced with Lentiviruses expressing Control-ShRNA or Kaiso-Specific ShRNA. 60 hours post transduction cells were trypsinized and seeded in cell culture treated plates and suspension plates (Ultra-low adherent plates) and cultured for 24 hours. Whole cell lysates were made from both the conditions and analysed for the indicated proteins by immunoblotting. Note that Kaiso is essential for repression of BMF only in cells cultured under suspension conditions but not in the adherent conditions. (B) Control and Kaiso-depleted cells as mentioned above from adherent plates and ultra-low adherent plates were reseeded in normal adherent plates and cultured for 24 hours more. Cell Viability was measured by crystal violet staining and fold change in absorbance was presented as bar diagram (bottom panel of B). Chromatin from MDA-MB 231 cells cultured in adherent plates as wells as ultra-low adherent plates was immunoprecipitated with Kaiso antibody and Normal mouse IgG. Kaiso binding to BMF promoter was analysed by PCR amplification of immunoprecipitated DNA using primers corresponding to a region of BMF promoter (−2139bp to −1943bp) harboring Kaiso Binding Sequence.

## Discussion

The transcription factor Kaiso has been shown to play tumor promoting role in solid cancers. However, its role in hematological cancers especially in MM has not been studied. Here, we documented an oncogenic addiction to Kaiso in MM and that several MM cell lines belonging to different MM genetic subgroups rely on Kaiso for survival regardless of the known oncogenic mutations harbored by these cell lines (Fig. 1). Mechanistically, we showed that Kaiso represses pro-apoptotic gene *bmf* by directly binding to its promoter at a region harboring the previously described Kaiso binding site (KBS) (58, 64), but not to a region lacking KBS (Fig. 2). Interestingly, while previous reports suggested that Kaiso binding to KBS in TGF-β receptor results in transcriptional activation leading to enhanced TGF-β signaling, our results indicated to that Kaiso binding to KBS in *bmf* promoter results in transcriptional repression than activation. This indicates that the transcriptional outcome (activation Vs repression) upon Kaiso binding to KBS may vary depending on the gene promoter and interacting partner/s. Here, we showed that in MM Kaiso interacts with HDAC1 to form Kaiso-HDAC1 complex (Fig. 3) and we could not observe Kaiso interaction with other HDACs in MM (data not shown). Importantly, our results indicated that HDAC1 is also essential to repress *bmf* in MM and that Kaiso is essential to recruit HDAC1 to *bmf* promoter to maintain deacetylated state as shown by the data that in Kaiso-depleted cells HDAC1 is not recruited to *bmf* promoter resulting in hyperacetylation of Histone H3 (Fig. 4). Moreover, exogenous HDAC1 overexpression was insufficient to repress *bmf* in Kaiso-depleted cells suggesting a clear dependence of HDAC1 on Kaiso for *bmf* repression (Fig. 4 and Supplementary Fig. 4). It is interesting to note that HDAC4 in a complex with RelB also was previously shown to repress *bmf* (75) and our results here identify HDAC1 also as a *bmf* repressor. It is however, unclear whether and how different HDACs coordinate to repress *bmf* and contribute to MM cell survival. Collectively, our data conclude that the Kaiso-HDAC1 complex maintains repressive chromatin on *bmf* promoter and thereby contribute to MM cell survival.

While the Kaiso-HDAC1 complex formation is necessary for *bmf* repression, it is not clear which oncogenic signal maintains high levels of Kaiso in MM cells. Although, TGF-β signaling was shown to regulate Kaiso expression in solid tumors (58), we did not observe a role for TGF-β in Kaiso regulation in MM (data not shown). However, we repeatedly observed that blocking the Jak-STAT pathway results in significantly reduced levels of Kaiso in MM cells (Fig. 5A). Among the STAT family of transcription factors, silencing STAT1 resulted in significantly reduced levels of Kaiso in MM cells (Fig. 5B). Interestingly, STAT1-depletion also resulted in highly elevated BMF levels like in the case of Kaiso-depletion resulting in MM cell death (Fig. 5B). Also, STAT1, depletion has completely abrogated subcutaneous MM tumor growth in nude mice (Fig. 5). While the role of STAT1 in MM has been confusing, data presented here provide conclusive evidence that STAT1 is absolutely essential for MM cell survival by regulating Kaiso expression and repressing *bmf*. Moreover, our data found that STAT1 is constitutively phosphorylated in most MM cell lines as well as in primary patient derived CD138+ MM cells (Fig. 6), further highlighting the oncogenic role for STAT1 in MM. Importantly, the data presented here identify Kaiso as a novel downstream effector for STAT1 in MM cell survival by repressing *bmf* as evidenced by the rescue of STAT1-depleted cells from apoptosis and reversal of *bmf* levels upon overexpression of exogenous Kaiso (Fig. 6 and Supplementary Fig. 6).

Although, our data revealed that STAT1 is constitutively phosphorylated in MM, what signal maintains constitutive STAT1 phosphorylation needs to be investigated further. Nevertheless, STAT1 is normally activated downstream of Type-1 or Type-II interferon signaling pathways and functions by forming a STAT2 – STAT1 heterodimer or a STAT1 homodimer respectively (78). Our results indicated that STAT2-depletion had no effect on MM cell survival and on *bmf* repression (Supplementary Fig. 5C and 5D) suggesting that STAT1 functions independent of STAT2 in MM.

Collectively, these data establish the STAT1-Kaiso-HDAC1-BMF axis as a novel survival mechanism operated in MM (Please see model in Fig.8). Thus, targeting the STAT1-regulation of Kaiso would offer new therapeutic strategy for MM. Importantly, as both Kaiso and HDAC1 are known to play crucial role/s in normal homeostasis and since Kaiso-HDAC1 complex is essential for *bmf* repression, development of strategies to disrupt the Kaiso-HDAC1 complex formation is expected to de-repress *bmf* and induce MM cell death and offer effective novel therapeutic strategy for MM with minimal or lesser toxic side effects. Such strategies would aid in MM cure either alone or in combination with currently approved MM therapy regime.

**Figure. 8:**
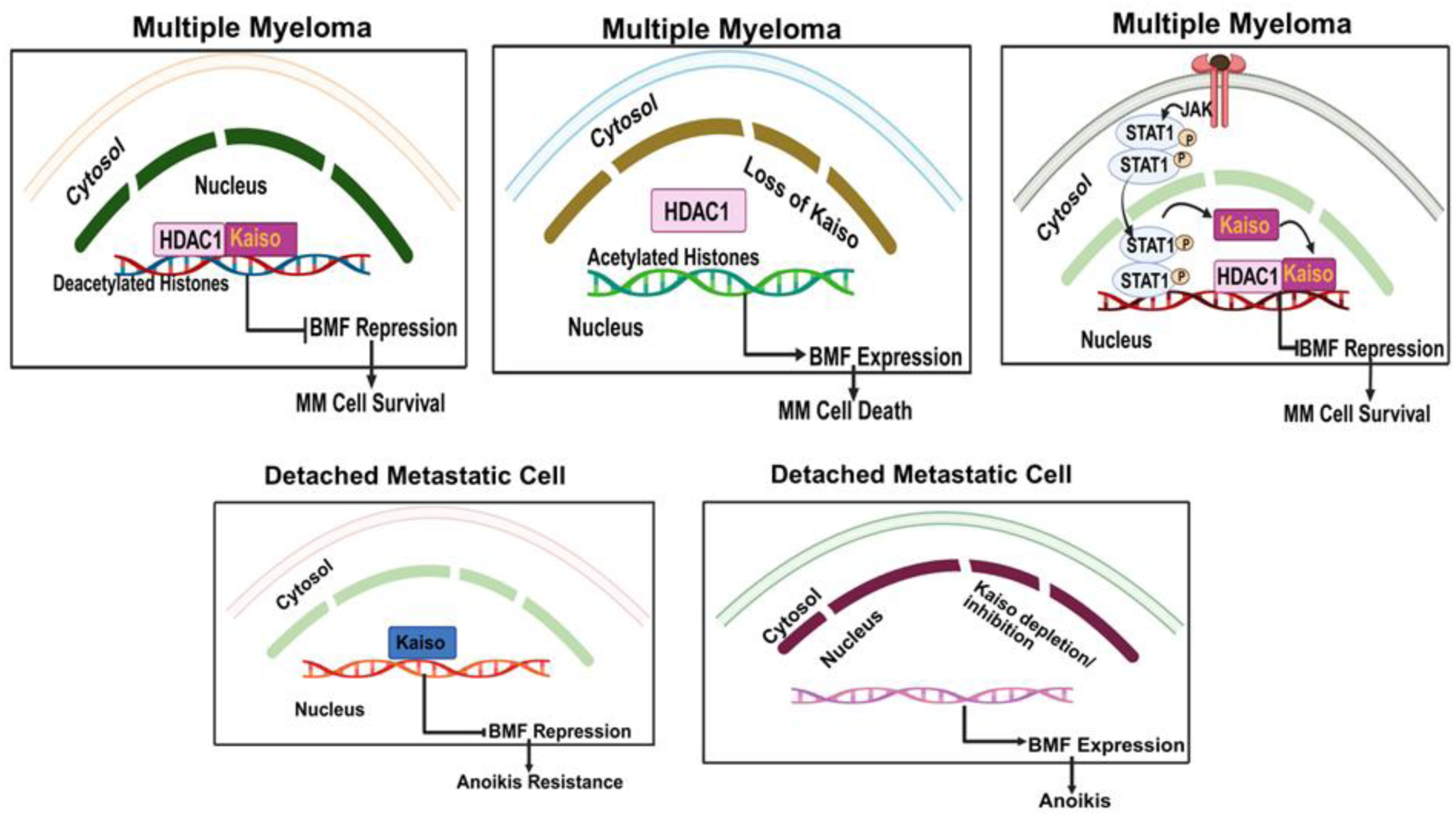
Model showing Kaiso dependent BMF repression in MM and metastatic solid tumors: **Kaiso-mediated *bmf* repression is a commonly operated survival mechanism in hematological malignancies (MM) and in Anoikis Resistance of Metastatic Solid Tumors. (A) Oncogenic addiction to Kaiso in MM**. Kaiso interacts with HDAC1 and recruits the same to *bmf* gene promoter to maintain repressive chromatin state and repress *bmf*. This helps in MM cell survival. **(B)** In the absence of Kaiso, HDAC1 is not recruited to *bmf* promoter resulting in active chromatin and *bmf* expression causing MM cell death. **(C) Kaiso is a novel downstream effector for oncogenic STAT1 in MM.** STAT1 regulates Kaiso expression, which in turn recruits HDAC1 to *bmf* promoter to repress the same causing MM cell survival. **(D) Kaiso is essential for anoikis resistance of metastatic solid tumors**. In metastatic solid tumors (Breast cancer, Glioma and Colorectal Cancer), Kaiso is recruited to *bmf* gene promoter. Loss of Kaiso does not affect *bmf* levels as long as the cells are adherent. However, Kaiso is absolutely essential to repress *bmf* gene in detached (essentially circulating) metastatic cancer cells suggesting a unique role of Kaiso in Anoikis Resistance.

The data presented here and other reports indicate that MM cells are very sensitive to BMF levels. But BMF is also known to be a key player in inducing anoikis death upon cellular detachment from extracellular matrix (ECM) (72–74). However, metastatic cancer cells exhibit anoikis resistance by repressing BMF (72–74). While, general repressors of BMF are known under both adherent and non-adherent conditions, whether BMF repression exclusively in detached but not in attached metastatic cancer cells is a key aspect of anoikis resistance has not been studied with mechanistic details. Our results indicate that Kaiso is essential exclusively in detached (non-adherent) but not in attached (adherent) metastatic breast cancer and glioma cells for *bmf* repression and survival of detached cells (Fig. 7). While these results establish Kaiso as a true regulator of anoikis resistance, a previous study suggested that in breast cancer Kaiso represses Wnt11 gene, whose function is essential for anoikis resistance. This finding is puzzling because Kaiso has previously been shown to promote metastasis but repression of Wnt11 gene by Kaiso suggests that Kaiso is a anoikis inducer and raises a question why would a metastasis promoter repress a gene essential for anoikis resistance. To this end, we silenced Kaiso in different metastatic cancers (Breast Cancer, Glioma and Colorectal Cancer) and found that it is an important contributor for anoikis resistance by repressing *bmf* (Please see model in Fig.8).

Collectively, our data revealed that Kaiso-mediated repression of *bmf* is a novel and commonly operated survival strategy in Multiple Myeloma and detached (non-adherent) metastatic solid tumor cells. Hence, targeting Kaiso-mediated *bmf* repression is expected to provide us with a platform to develop broad-spectrum cancer drug.

---

**Supplementary Fig. 1:**
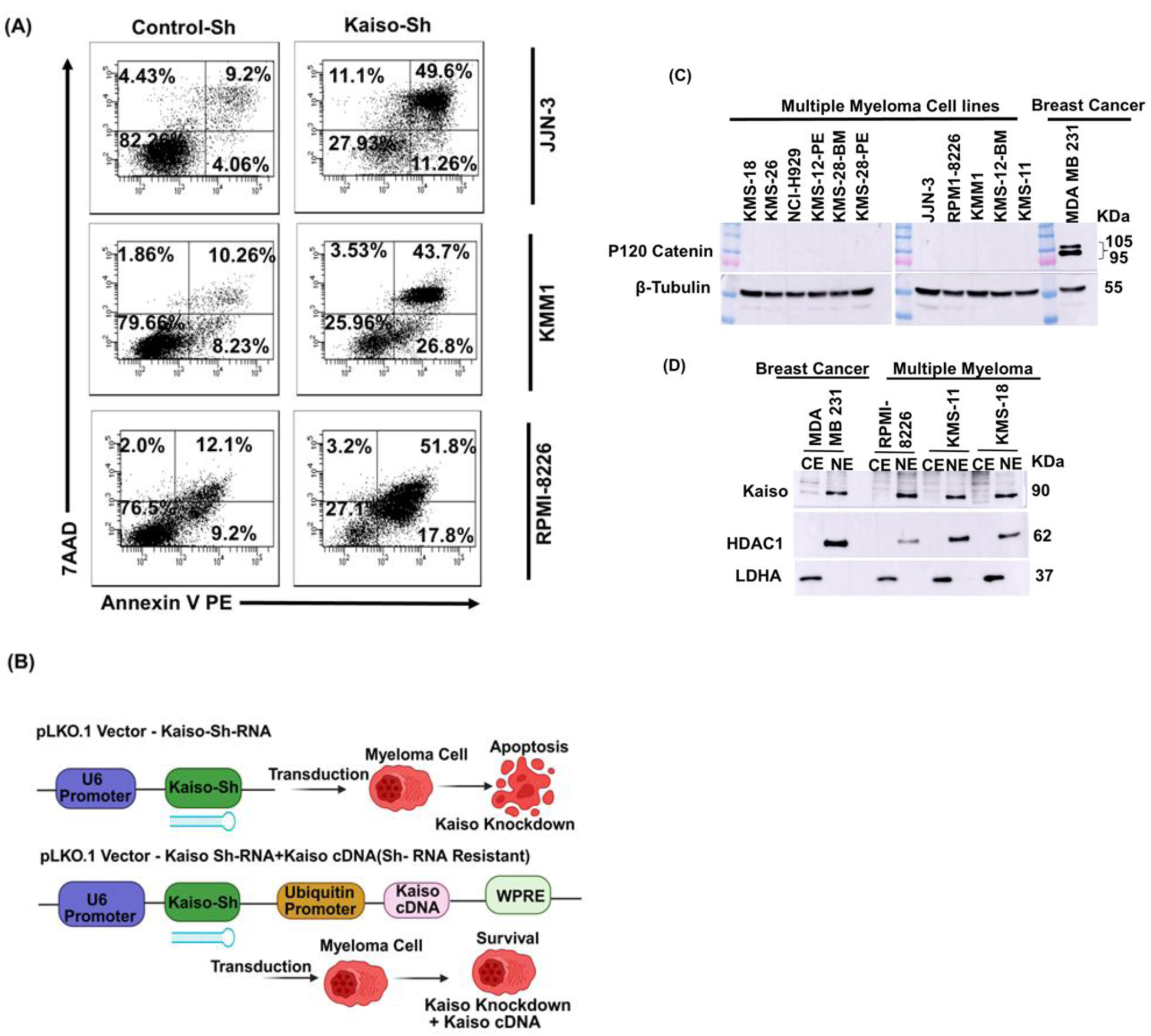
(A) Indicated MM cell lines were transduced with lentiviruses expressing control-ShRNA or Kaiso-ShRNA. 5 days post infection, cell viability was analysed by flow-cytometry upon staining with Annexin-V-PE and 7AAD. (B) Schematic model for Kaiso rescue construct design. The expression cassette of Kaiso cDNA consisting of ubiquitin promoter and wood chuck regulatory elements was cloned into pLKO.1 vector that expresses Kaiso-ShRNA. (C) Whole cell lysates from the indicated MM cell lines and breast cancer cells were analysed for p120 catenin expression by immunoblotting. Note that its expression is not detected in MM cell lines unlike MDA-MB 231 where it is abundantly expressed. (D) Nuclear and Cytoplasmic extracts from the indicated MM cell lines and breast cancer (MDA-MB-231) cells were analysed by immunoblotting analysis for the indicated proteins. Note that Kaiso is largely confined to Nucleus unlike MDA-MB 231.

**Supplementary Figure 2:**
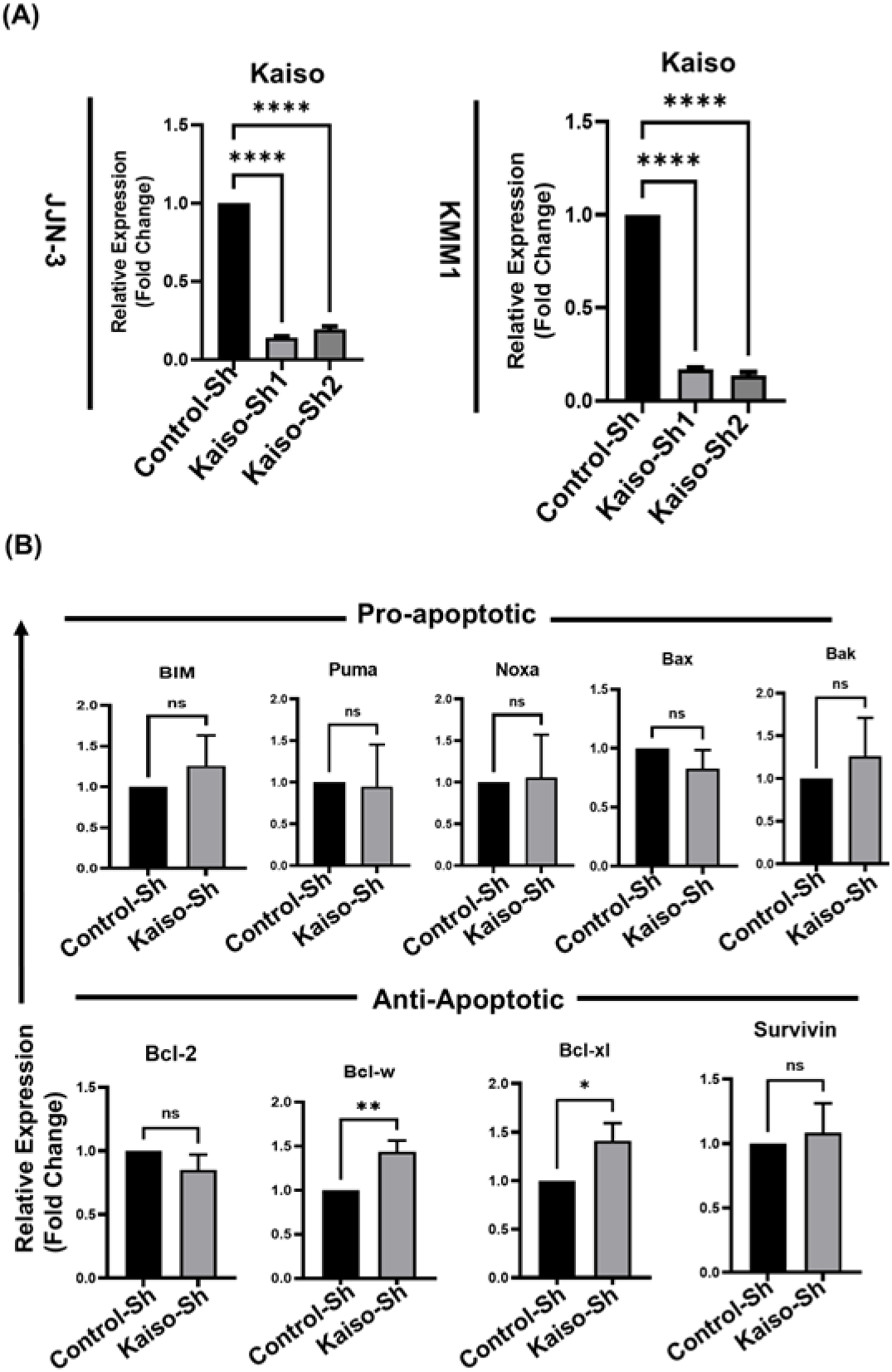
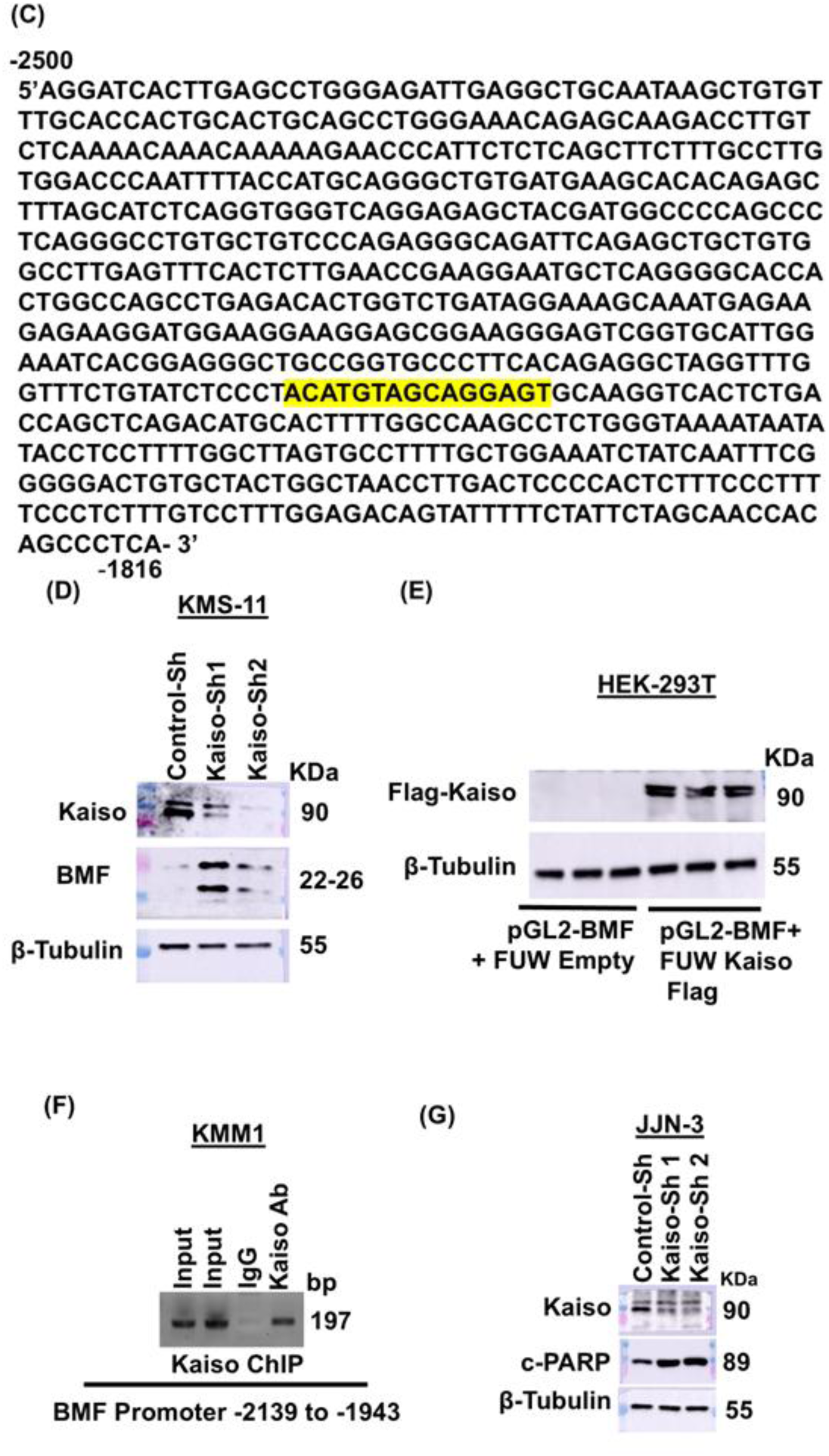
(A) JJN-3 and KMM1 cells were transduced with lentiviruses expressing Control-ShRNA or Kaiso-ShRNA. 60 hours post infection total RNA was made and analysed Kaiso knockdown by RT-PCR. (B) Total RNA from control and Kaiso-depleted JJN3 cells was analyzed by RT-PCR for the indicated pro- and anti-apoptotic genes. (C) Bio-informatic analysis of 2.5Kb BMF promoter by Mat-inspector for identifying potential transcription factor binding sites. Note that BMF promoter has strong Kaiso Binding sequence at (−2035bp to −2027bp) (KBS-TCCTGCNA / TNGCAGGA). BMF promoter sequence was obtained from UCSC genome browser. (D) KMS-11 cells infected with lentiviruses expressing Control-ShRNA or Kaiso-ShRNA. 60 hours post infection whole cell lysates were made and analysed for the indicated proteins by immunoblotting. (E) Immunoblotting analysis of the lysates used for BMF-promoter luciferase reporter assay (Fig.2C) for the indicated proteins. (F) Chromatin from KMM1 cells was immunoprecipitated with anti-Kaiso antibody or normal mouse-IgG. Kaiso binding to BMF promoter was analyzed by PCR amplification of chromatin-immunoprecipitated DNA using primers corresponding to a region of BMF promoter (−2139 to −1943) harboring Kaiso binding sequence. (G) Whole cell lysates from JJN-3 cells infected with lentiviruses expressing Control-ShRNA or Kaiso-ShRNA and analysed for the expression of indicated proteins by immunoblotting. Note that kaiso depletion significantly increased the c-PARP levels.

**Supplementary Figure. 3:**
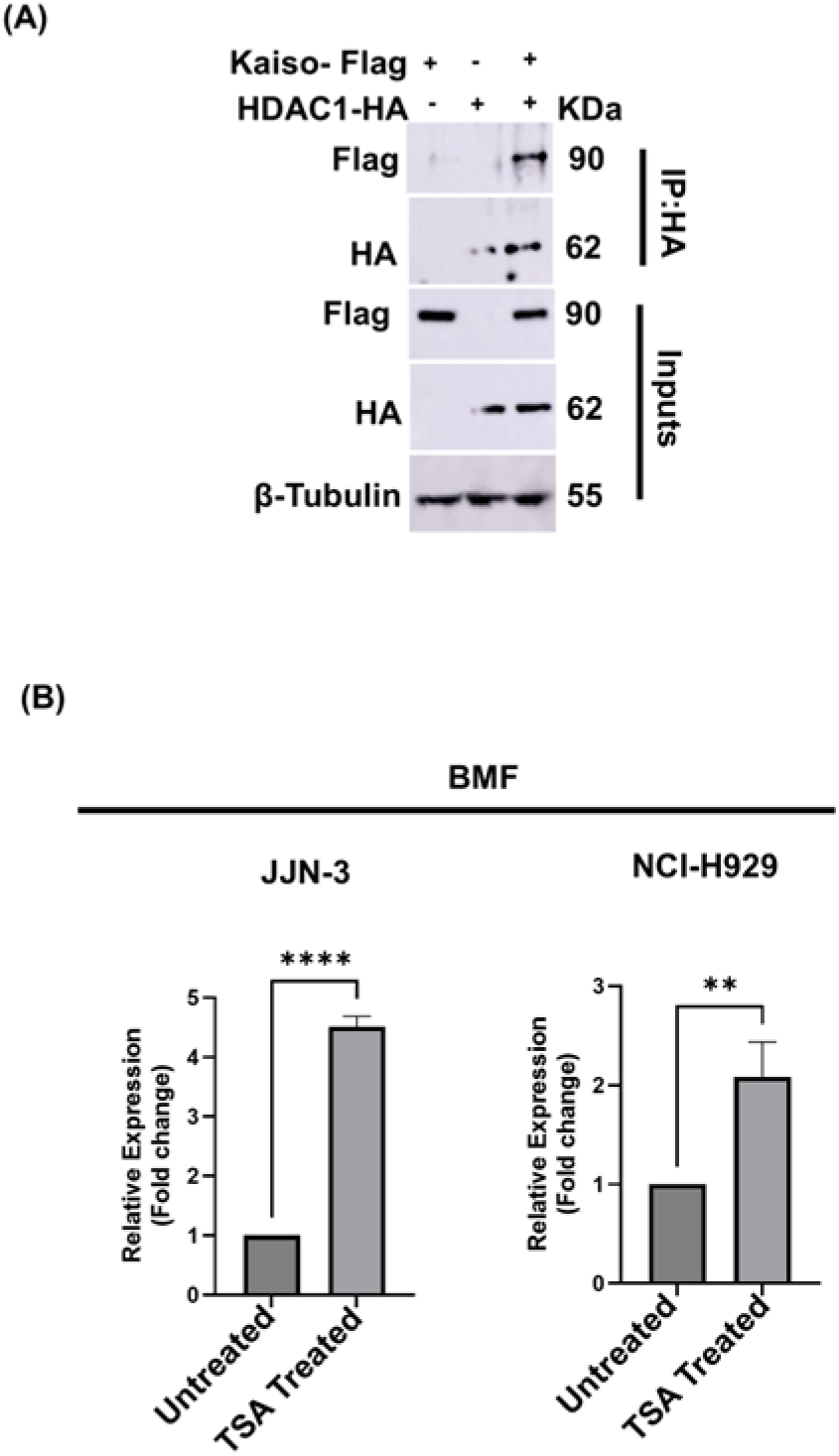
(A) HEK-293T cells were co-transfected with Kaiso-Flag and HDAC1-HA and immunoprecipitation was performed with anti-HA antibody and HDAC1-Kaiso interaction was analysed by immunoblotting as indicated. (B) Indicated Myeloma cells were treated with Pan HDAC inhibitor Trichostatin-A (TSA) for 2 hours. Total RNA was made and analysed for the relative expression of BMF by RT-PCR. Total RNA from DMSO treated cells was used as control.

**Supplementary Figure 4:**
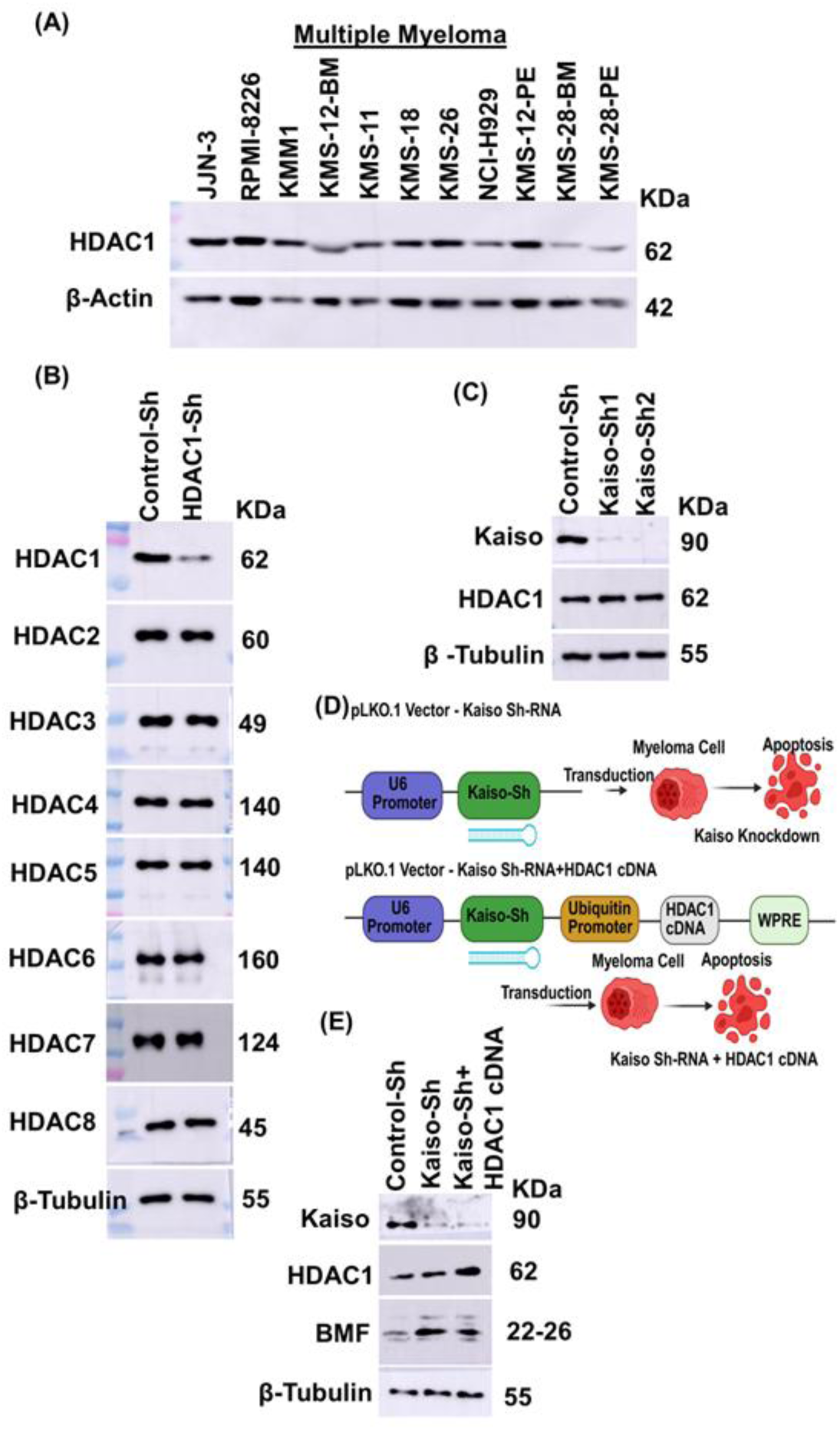
(A) Whole cell lysates from indicated multiple myeloma cells were analysed for the expression of HDAC1 levels by immunoblotting. Note that HDAC1 expression is abundantly expressed in most MM cell lines. (B) Whole cell lysates from control and HDAC1-depleted JJN3 cells were analyzed by immunoblotting for the indicated proteins. Note that HDAC1 depletion did not affect the expression of other HDACs. (C) Whole cell lysates from control and Kaiso-depleted JJN3 cells were analysed by immunoblotting for the indicated proteins. Note that HDAC1 protein levels were unaffected in Kaiso depleted cells. (D) Model for a lentiviral vector that co-expresses Kaiso-ShRNA and HDAC1-cDNA. HDAC1 cDNA expression cassette comprising ubiquitin promoter, HDAC1-cDNA and Wood Chuck regulatory elements was cloned into pLKO.1 Vector expressing Kaiso-ShRNA. (E) JJN-3 cells were transduced with Lentiviruses expressing Control-ShRNA, Kaiso-ShRNA and Kaiso-ShRNA+HDAC1 cDNA. 60 hours post infection whole cell lysates were made and analysed for indicated proteins by immunoblotting.

**Supplementary Fig. 5:**
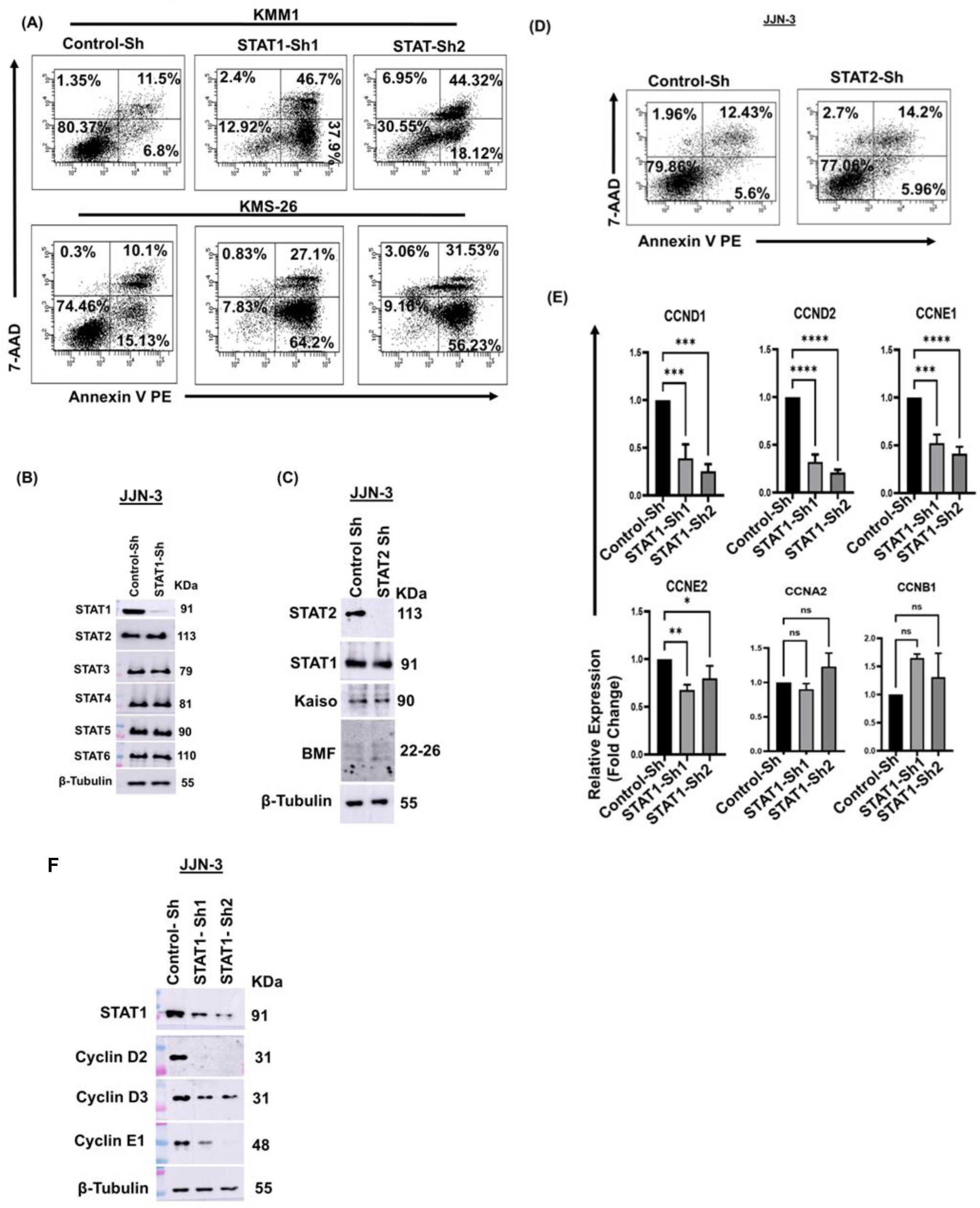
(A) KMM-1 and KMS-26 cells were infected with lentivirus expressing Control-ShRNA or STAT1-ShRNA. 5 days post infection cell viability was analysed by flow-cytometry upon staining with Annexin V-PE and 7-AAD. (B) Whole cell lysates from control and STAT1-depleted JJN3 cells were analysed for the indicated proteins by immunoblotting. Note that STAT1 depletion did not affect the expression of other STATs indicating the specificity of STAT1-ShRNA. (C) Whole cell lysates from control or STAT2-depleted JJN-3 cells were analysed by immunoblotting for the indicated proteins. (D) JJN3 cells were transduced by lentiviruses expressing Control-ShRNA or STAT2-ShRNA. 5 days post infection cell viability was analysed by flow-cytometry upon staining with Annexin V-PE and 7-AAD. Note that STAT2-depletion had no effect on Kaiso expression and MM cell viability. (E) Total RNA and (F) whole cell lysates were made from control and STAT1-depleted JJN-3 cells and analysed by RT-PCR for the indicated genes and by immnoblotting for the indicated proteins.

**Supplementary Figure 6:**
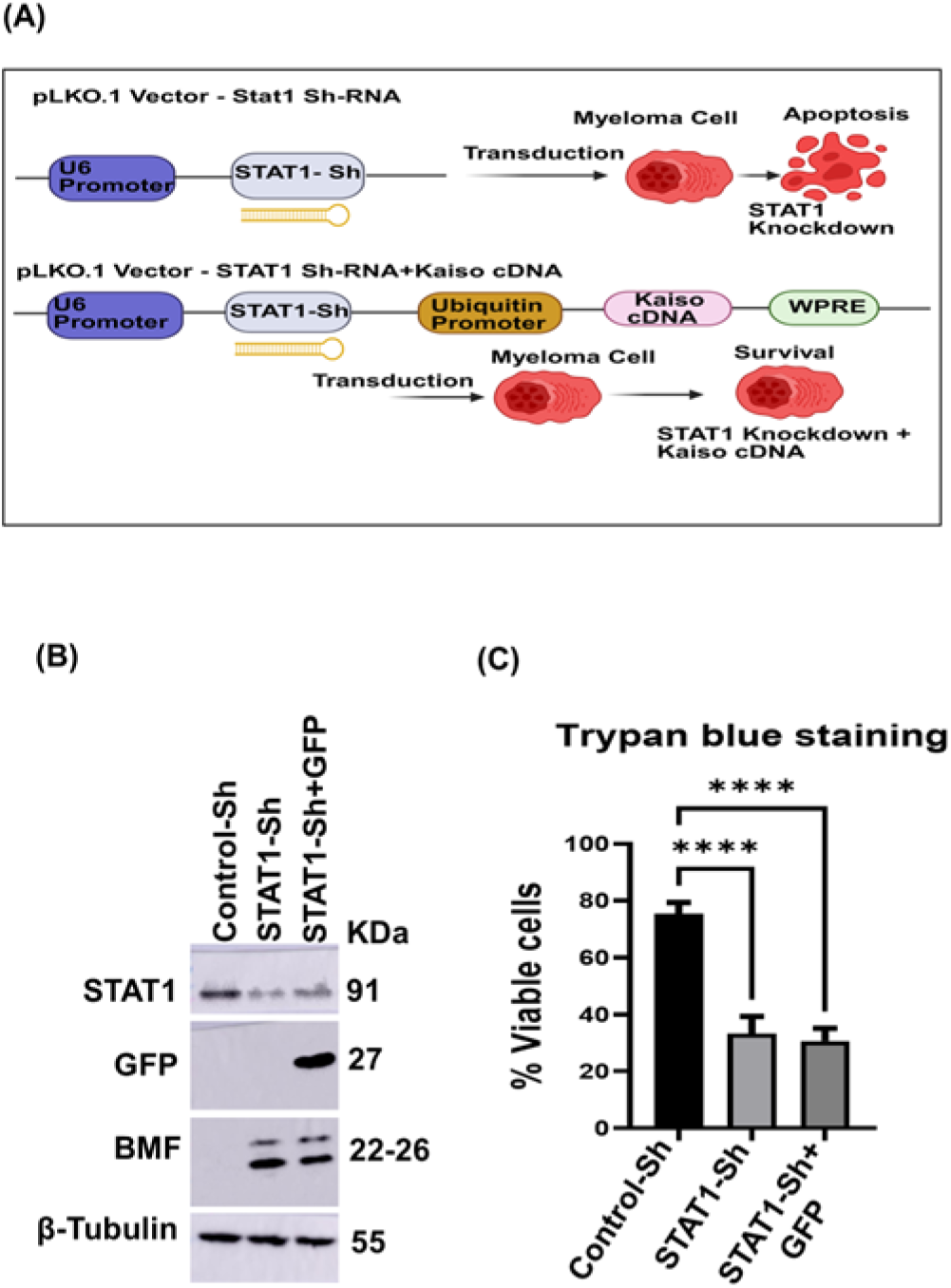
(A) Model for lentiviral vector that co-expresses STAT1-ShRNA and Kaiso cDNA. Kaiso expression cassette comprising ubiquitin promoter, Kaiso-cDNA and Wood Chuck regulatory elements was cloned into pLKO.1 expressing STAT1-ShRNA. (B) EGFP was over expressed in STAT1-depleted JJN-3 cells in a manner similar to Kaiso-overexpression as described in (A). Whole cell lysates were prepared from control cells, STAT1-depleted cells and STAT1-depleted EGFP expressing cells and were analysed for the levels of indicated proteins by immunoblotting. (C) JJN3 cells were transduced with lentiviruses expressing control-ShRNA or STAT1-ShRNA or STAT1-ShRNA + EGFP cDNA. 5 days post infection, cell viability by was measured by Trypan blue statining. Note that EGFP overexpression did not impact BMF expression and could not rescue STAT1 depleted cells from apoptosis.

**Supplementary Figure 7:**
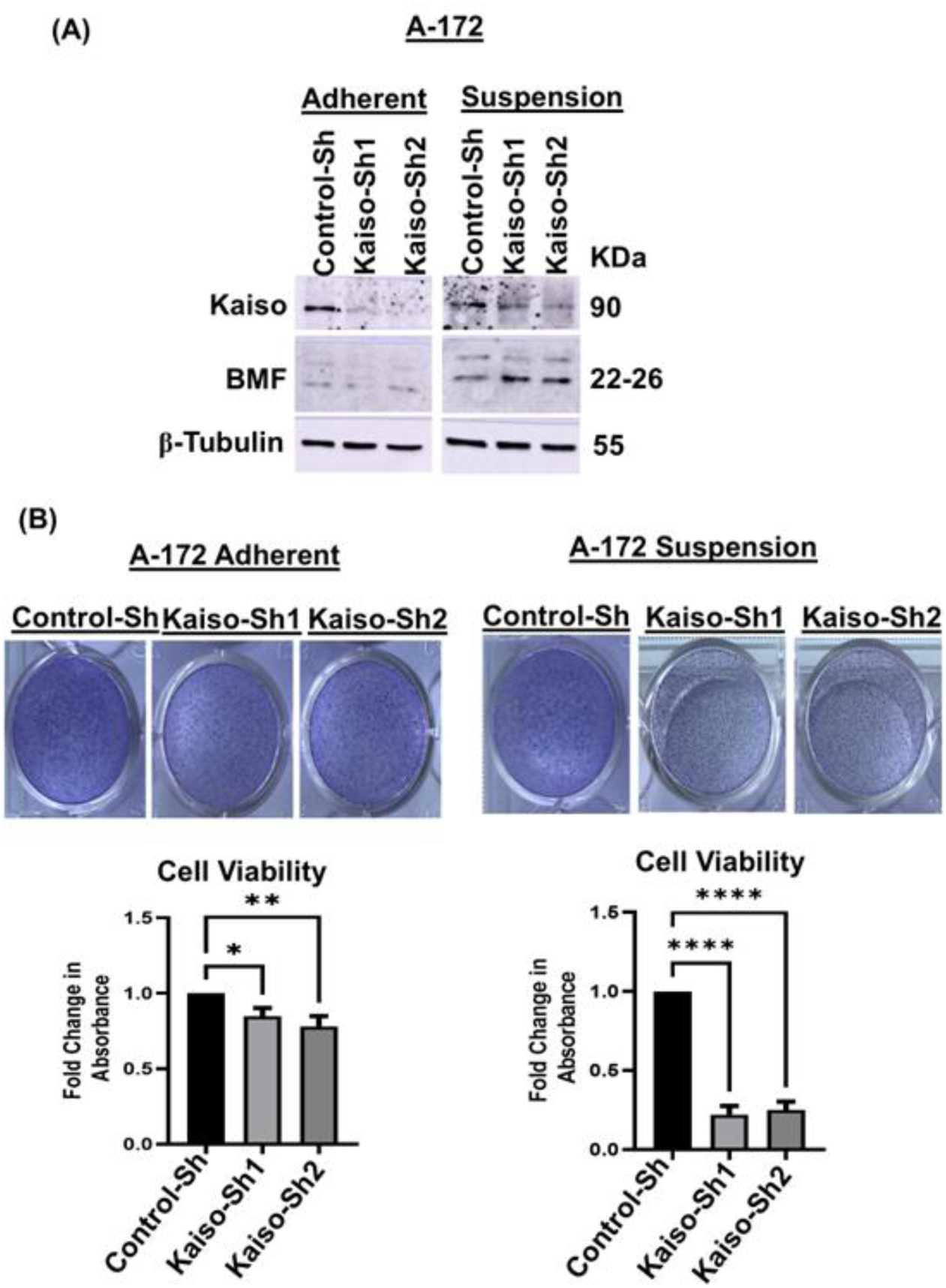
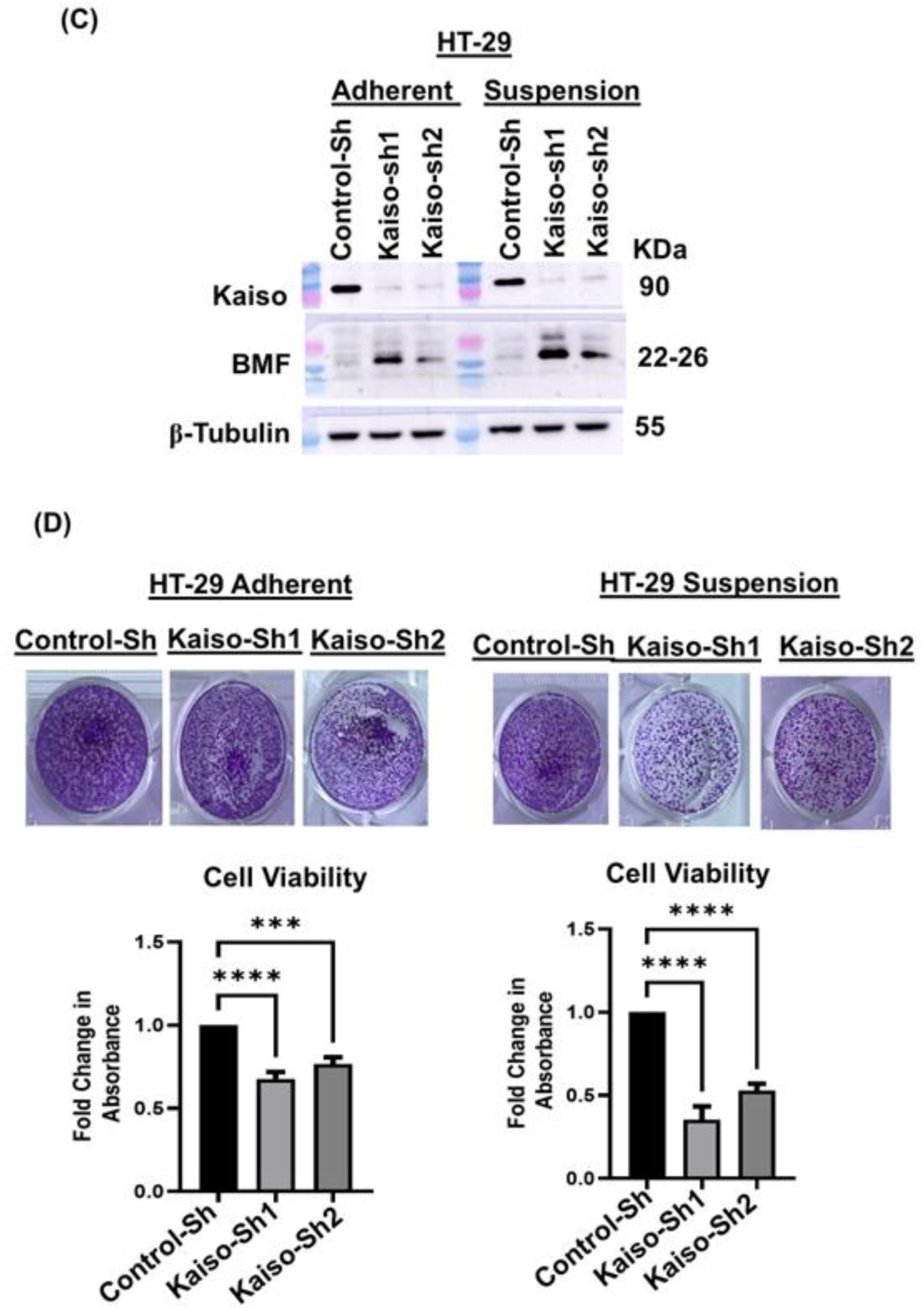
Kaiso is essential for Anoikis resistance in Glioma and Colorectal Cancer. (A) A-172 (Glioblastoma) cell line was transduced with Lentiviruses expressing Control-ShRNA or Kaiso-Specific ShRNA. 60 hours post transduction cells were trypsinized and seeded in cell culture treated plates (adherent) and suspension plates (Ultra-low adherent plates) and cultured for 24 hours. Whole cell lysates were made from both the conditions and analysed for the indicated proteins by immunoblotting. Note that Kaiso is essential for repression of BMF only in cells cultured under suspension conditions but not in the adherent conditions. (B) Control and Kaiso-depleted cells as mentioned above from adherent plates and ultra-low adherent plates were reseeded in adherent plates and cultured for 24 hours more. Cell Viability was measured by crystal violet staining fold change in absorbance was presented as bar diagram (bottom panel of B). (C) HT-29 (Colorectal cancer) cell line was transduced with Lentiviruses expressing either Control-ShRNA or Kaiso-Specific ShRNA and were cultured under similar conditions as explained above for glioma cells and immunoblotting analysis was performed for the indicated proteins. (D) Control and Kaiso-depleted HT-29 cells were cultured in a similar way as explained above for glioma cells and cell viability was analysed by crystal violet staining. Bar diagram (bottom pannel) indicates fold change in relative absorption in crystal violet assay. Note that Kaiso-depletion results in elevation of BMF levels predominantly in detached cells compared to adherent cells and kaiso is essential for survival of detached cells but not adherent cells.

## Acknowledgements

This work was supported by Wellcome Trust – DBT India Alliance intermediate fellowship to SV. RM was supported by CSIR fellowship. All the MMCLs used in this study were kindly provided by Prof. Leif Bergsagel and Prof. Marta Chesi. We sincerely thank Prof. Rashna Bhandari, Dr. Pranjali Pore, Mr. Pavan and Mr. Bheemudu, Tanmay Kumar Mohanty and Abhijeet Behera for helping us with xenograft experiments at the animal facility, CDFD, Hyderabad, India.

## Author Contributions

Sivakumar Vallabhapurapu has designed the whole work, conducted the research, wrote the manuscript and obtained funding for this research. Ramadevi Mutra has designed and conducted the experiments and wrote the manuscript. Faiha Mundodan, Prajakta Ghatage and Asha Vallem have helped with conducting few western blots. Anciya VR participated in cloning and western blots. Charles Lawrie, Carla Sole, and Carlos Panizo provided primary MM cell lysates.

